# Degradation of ribosome and chaperone proteins is attenuated during the Differentiation of Replicatively Aged C2C12 Myoblasts

**DOI:** 10.1101/2021.10.18.464830

**Authors:** Alexander D. Brown, Claire E. Stewart, Jatin G. Burniston

**Affiliations:** Research Institute for Sport & Exercise Sciences, Liverpool John Moores University, Liverpool, UK

**Keywords:** Proteomics, Aging, Muscle Regeneration, Protein Synthesis, Protein Degradation

## Abstract

Age-related impairments in myoblast differentiation may contribute to reductions in muscle function in older adults, however, the underlying proteostasis processes are not well understood. Young (P6-10) and replicatively aged (P48-50) C_2_C_12_ myoblast cultures were investigated during early (0h-24h) and late (72h-96h) stages of differentiation using deuterium oxide (D_2_O) labelling and mass spectrometry. The absolute dynamic profiling technique for proteomics (Proteo-ADPT) was applied to quantify the absolute rates of abundance change, synthesis and degradation of individual proteins. Proteo-ADPT encompassed 116 proteins and 74 proteins exhibited significantly (P<0.05, FDR <5 %) different changes in abundance between young and aged cells at early and later periods of differentiation. Young cells exhibited a steady pattern of growth, protein accretion and fusion, whereas aged cells failed to gain protein mass or undergo fusion during later differentiation. Maturation of the proteome was retarded in aged myoblasts at the onset of differentiation, but their proteome appeared to ‘catch up’ with the young cells during the early phase of the differentiation period. However, this ‘catch up’ process in aged cells was not accomplished by higher levels of protein synthesis. Instead, a lower level of protein degradation in aged cells was responsible for the elevated gains in protein abundance. Our novel data point to a loss of proteome quality as a precursor to the lack of fusion of aged myoblasts and highlights dysregulation of protein degradation, particularly of ribosomal and chaperone proteins, as a key mechanism that may contribute to age-related declines in the capacity of myoblasts to undergo differentiation.

## Introduction

Skeletal muscle is the most abundant tissue in young healthy adults and is essential for the maintenance of posture and the activities of daily living. Adult muscle consists of terminally differentiated myofibres and relies on muscle satellite cells to regenerate [1]. Older adults experience losses in muscle size and physical strength that impinge on their independence, increase their risk of falling and reduce their quality-of-life. Indeed, beyond the age of 60 years, humans may experience an ~30 % decline in muscle mass [2,3] and the rate of loss of muscle can be exacerbated by low levels of physical activity or periods of bedrest associated with hospitalisation or illness. The capability of muscle to regenerate following exercise or injury also declines with advancing age and further contributes to the reductions in physiological function and muscle homeostasis [4]. Muscle regeneration is dependent on the ability of myoblasts (activated satellite cells), to undergo terminal differentiation and fuse to form mature myofibers *in vivo*, or myotubes *in vitro*. Myoblasts generated from the muscle of older humans exhibit compromised fusion [5], and the loss of functionality, in addition to decreases in satellite number, underlies the age-related decline in muscle regenerative capacity [4].

Losses in muscle regenerative capacity may be a consequence of factors that are either intrinsic or extrinsic to muscle cells [6]. Parabiotic studies conducted *in vivo* [7] and *in vitro* [8,9] have established an important role of the host environment on muscle regeneration and satellite cell function, whilst replicative aging of cells *in vitro* has demonstrated age-related impairments in differentiation capacity [10] that occur in the absence of the confounding effects of changes in systemic environment. Replicatively aged C_2_C_12_ myoblasts exhibit impaired differentiation and fusion [11,12] similar to human replicatively-aged primary myoblasts [13] and cells and tissue isolated from older rodents and humans [13–18]. Myoblast differentiation is orchestrated by complex programmes of cell signalling [19] and altered gene expression [17] that result in dynamic changes to the cell proteome during differentiation [20,21]. Accordingly, the reductions in differentiation and fusion capacity of replicatively aged cells are accompanied by impaired cell signalling and gene expression [22,23]. Insulin-like growth factors I/II (IGF-I and II) are critical for cell proliferation, differentiation and growth [24] and are key regulators of muscle hypertrophy. Following activation of the IGF-I receptor, phosphorylation and activation of downstream signalling molecules, including protein kinase B (Akt) and mechanistic target of rapamycin (mTOR) increase protein translation and ribosomal biogenesis, and thus myotube hypertrophy [19,25]. Age-related reductions in Akt activity affect muscle mass by directly reducing transcription of myogenic regulator factors (MRFS) MyoD and myogenin, which negatively impact myoblast proliferation, differentiation and fusion [17,24,26]. In addition, the inflammatory protein tumour necrosis factor alpha (TNF- α) and myostatin which is a member of the transforming growth factor-β superfamily and a negative regulator of muscle hypertrophy [27] are both increased with ageing and each impairs myoblast differentiation [26,28]. Elevations in TNF-α during ageing contribute to impaired myoblast differentiation through reductions in mitogen-activated protein kinase (MAPK) p38α and thus myocyte enhancer factor-2 (MEF2) signalling [12,29,30] as well as through the activation of nuclear factor kappa β (NFKβ) which culminates in protein loss in myotubes [31].

Currently, there is limited proteomic data regarding the effect of ageing on myoblast differentiation. Comprehensive age-related changes to the proteome of mature human muscle have been reported [32,33], including reductions in contractile, mitochondrial, glycolytic, ribosomal, anti-inflammatory and proteostasis proteins with advancing age. On the other hand, pro-inflammatory, spliceosome and DNA damage recognition and repair proteins become more abundant in the muscle of older humans [32,33]. These findings agree with studies in aged flies [34], worms [35] and the muscle of older laboratory mice [36], which also report a lesser abundance of ribosomal, glycolytic and proteostasis proteins as a function of age. A loss in proteostasis is an acknowledged hallmark of ageing [37] that is conserved across model organisms [34,35], laboratory species [36,38,39] and humans [32,33,40,41]. Protein turnover is an important component in the maintenance of proteome quality, i.e. proteostasis, in addition to changes to the abundance and activity of protein chaperones. Studies in flies [34], worms [35], mice [38] and humans [42] have consistently reported age-related declines in protein turnover in adult muscle. The aforementioned changes to the abundance profile of proteins in human muscle are also accompanied by a decline in protein turnover [40,41,43] as well as dampened responses to anabolic stimuli such as feeding and exercise [44] in the muscle of older humans. Proteomic studies reporting protein turnover in fast-twitch extensor digitorum longus (EDL) and slow-twitch soleus of aged laboratory mice [38] found that advancing age is associated with a greater turnover and abundance of mitochondrial proteins specifically in fast-twitch muscle but respiratory function was depressed and the generation of hydrogen peroxide by Complex I was elevated in fast-twitch muscle of older mice [38], indicating that dysfunctional proteins may accumulate in older muscle.

During ageing the accumulation of oxidatively damaged and misfolded proteins may overwhelm chaperone systems and exceed the capacity for protein turnover to replace damaged proteins leading to losses in cellular function [45]. A collapse of proteostasis is an acknowledged component of skeletal muscle ageing [46] and, compared to other tissues, muscle fibres may experience additional burdens on proteostasis specifically related to mechanical stresses [47,48]. Protein degradation is the crucial cellular process that removes damaged or misfolded proteins and, thereby, helps to prevent proteostasis collapse and the accumulation of protein aggregates that may become toxic to the cell [49]. Degradation of proteins is mediated by the ubiquitin proteasome system (UPS) or autophagy processes and, in the majority, research has focused on age-related changes to UPS [46]. Stimulation of UPS can extend lifespan in C. elegans [50] but few studies have specifically investigated changes to the balance between protein synthesis and protein degradation that, together, constitute protein turnover. We have sought to address this limitation by developing dynamic proteomic profiling methods [51] that simultaneously acquire data on both the abundance and synthesis rate of individual proteins.

The measurement of protein abundance alongside protein-specific synthesis rates affords new opportunities to study protein dynamics in systems that are undergoing change [52]. Furthermore, our Absolute Dynamic Profiling Technique for Proteomics (Proteo-ADPT), is capable of generating data in mole and absolute (ng) units, which benefit the biological interpretation of turnover data [21]. Changes to the abundance of each protein across different time points is used to calculate the absolute rate of abundance change, which is driven by changes to the absolute synthesis rates (ASR) and/ or the absolute degradation rates (ADR) on a protein-by-protein basis [52]. When measured at the level of individual proteins, changes in protein abundance that are not matched by equivalent changes to the synthetic rate of the protein may be attributed to protein degradation. In the current work we have applied this technique to study the contributions of protein synthesis and degradation to changes in protein abundance that occur during early and later stages of differentiation in young and replicatively aged murine C_2_C_12_ cells.

## Methods

### Cell Culture and Deuterium Oxide Labelling

All cell culture experiments were performed under sterile conditions and the incubation environment was standardised at 5 % CO_2_ at 37 °C (HERAcell 150i incubator; Thermo Scientific). C_2_C_12_ murine myoblasts were purchased from ATCC (Rockville, MD, USA) and divided into two groups: (1) young (low passage 6 - 10) and (2) replicatively aged (high passage 48 - 50). The aged C_2_C_12_ myoblasts were based on a previous model of replicative aging [11,12]. In brief, the aged myoblasts underwent 140 - 150 population doublings. For all experimental conditions, cells were seeded (1 x 10^5^ cells/ml) on gelatinised 6 well plates (Nunc, Roskilde, Denmark) and grown to ~80 % confluency. When confluent, growth media (GM: Dulbecco’s Modified Eagle Medium (DMEM), 10 % heat-inactivated fetal bovine serum (FBS), 10 % heat-inactivated new-born calf serum, 2 mM L-glutamine, and 1 % penicillin-streptomycin) was removed, cells were washed twice with phosphate buffered saline (PBS) and 2 ml differentiation media (DM: DMEM, 2 % heat-inactivated horse serum, 2 mM L-glutamine, 1 % penicillin-streptomycin) was added in the absence or presence of D_2_O. The protocol was designed to investigate the impact of age on early and late differentiation, therefore the first experimental protocol involved culturing the myoblasts in DM +/− D_2_O from 0 h - 24 h (Early Differentiation). The second protocol involved culturing the myoblasts in DM for 72 h, enabling the formation of myotubes (Late Differentiation). The 72 h timepoint was considered the beginning of the second experimental protocol and the myotubes were transferred to fresh DM +/− D_2_O for a further 24 h (72 h - 96 h). Isotopic labelling of newly synthesised proteins was used to quantify the fractional synthetic rates by supplementing the 2 ml DM with 99.8 % D_2_O (4 % enrichment) from 0 h - 24 h and from 72 h - 96 h. Experimental controls were treated equivalently in the absence of D_2_O. Cells were extracted in the absence or presence of D_2_O at 0 h, 24 h, 72 h and 96 h.

### Protein Extraction and Quantification

Following treatments and timings detailed above, cell imaging was performed prior to DM aspiration and monolayer washing, twice with ice cold PBS. Cells were lysed with 250 μl of 1x RIPA buffer (Sigma-Aldrich, Poole, UK) for 5 min at 4 °C, scraped into Eppendorf tubes in preparation for total protein quantification, digestion and proteomic analyses. Total protein concentration (μg/μl) was quantified using the BCA™ protein kit (Rockford, IL, USA). BCA standards were prepared by serial dilution of bovine serum albumin (BSA) from 2 mg/ml to 0 mg/ml. For analyses, 20 μl of standard or sample was added to each well. All the samples were pipetted in duplicate prior to 200 μl working agent being added to each well, according to manufacturer instructions. The samples were incubated at 37 °C for 60 min and scanned at an absorbance of 595 nM. Protein concentrations of cell samples were interpolated from the standard curve by linear regression.

### Muscle Cell Digestion and Dilution

Proteins were digested and diluted consistent with our previous work [52]. Lysates containing 100 μg protein were precipitated in 5 volumes acetone for 1 h at −20 °C and the protein pellets were resuspended in 200 μl of UA buffer (8 M urea, 100 mM tris, pH 8.5). Samples were incubated at 37 °C for 15 min in UA buffer with 100 mM dithiothreitol (DTT) followed by 20 min at 4 °C in UA buffer containing 50 mM iodoacetamide (protected from light). Samples were washed twice with 100 μl UA buffer and transferred to 50 mM ammonium hydrogen bicarbonate (Ambic). Sequencing grade trypsin (Promega; Madison, WI, USA) in 50 mM Ambic was added at an enzyme to protein ratio of 1:50 and the samples were digested overnight at 37 °C. To terminate digestion, peptides were collected in 50 mM Ambic and trifluoracetic acid (TFA) was added to a final concentration of 0.2 % (v/v). Aliquots, containing 4 μg peptides, were desalted using C18 Zip-tips (Millipore, Billercia, MA, USA) and eluted in 50:50 of acetonitrile and 0.1 % TFA. Peptide solutions were dried by vacuum centrifugation for 25 min at 60 °C and peptides were resuspended in 0.1 % formic acid spiked with 10 fmol/ul yeast ADH1 (Waters Corp.) in preparation for LC-MS/MS analysis.

### Liquid Chromatography-Mass Spectrometry

Peptide mixtures were analysed by nanoscale reverse-phase ultra-performance liquid chromatography (NanoAcquity; Waters Corp.) and online electrospray ionisation quadrupole time-of-flight mass spectrometry (QToF Premier; Waters Corp.). Samples were loaded on to a Symmetry C18 5 μm, 2 cm x 180 μm trap column (Waters, Milford, MA) in 2.5 % ACN, 0.1 % (v/v) formic acid. Separation was conducted at 35 °C through a BEH C18 1.7 μm, 25 cm x 75 μm analytical reverse phase column (Waters, Milford, MA) using a linear gradient from 2.5 % to 37.5 % acetonitrile in 0.1 % (v/v) formic acid over 60 min at a flow rate of 300 nL/min. The mass spectrometer was operated in a data-dependent positive electrospray ionisation mode at a resolution of >10,000 full width at half maximum. Prior to sample analysis, the time-of-flight analyser was calibrated using fragment ions of Glu-1-fibrinopeptide B from 50 to 1990 m/z. LC-MS profiling recorded peptides between 350 m/z and 1500 m/z using MS survey scans of 0.9 s duration with an inter-scan delay of 0.1 s. Equivalent data-dependent tandem mass spectrometry (MS/MS) spectra were collected from the control samples with no D_2_O label. MS/MS spectra of fragment ions were recorded over 50-2000 m/z for the 5 most abundant precursors ions of charge 2+ or 3+ detected in the survey scan. Precursor fragmentation was achieved by collision induced dissociation at an elevated (20-40 eV) collision energy over a duration of 0.25 s scan an inter-scan delay of 0.05 s. A dynamic exclusion window was set to 30 s to avoid repeat selection of prominent ions. Acquisition was switched from MS to MS/MS mode when the base peak intensity exceeded a threshold of 30 counts. MS mode was restored when the total Ion Chromatogram in the MS/MS channel exceeded 7500 counts/s or when 1 s (5 scans) were acquired.

### Proteome profiling

Protein abundance profiles were analysed using Progenesis Quantitative Informatics for Proteomics (QI-P; Nonlinear Dynamics, Newcastle, UK). Spectra were aligned, using prominent ion features (mean ± SD per chromatogram: 655 ± 42) to a reference chromatogram. Each chromatogram analysis window encompassed 15-105 min and 350-1500 m/z and 32,824 features were identified with charge states of +2, +3 or +4. Log transformed MS data (peptide abundance) was normalised by inter sample abundance ratio. MS/MS spectra in Mascot generic format and peptide sequences were searched against Swiss-Prot database. The search was restricted to ‘mus musculus’ which covered 17,006 sequences and performed using a locally employed Mascot server (www.matrixscience.com; version 2.2.03). The enzyme specificity was as follows: trypsin allowing for 1 missed cleavage, carbamidomethyl modification of cysteine (fixed), oxidation of methionine (variable) and deamidation of asparagine and glutamine (variable). The m/z error was set at ± 0.3 Da. The false discovery rate (FDR) based on decoy database search was 1.88 %. The output file (Mascot; xml format) was restricted to nonhomologous protein identification. Before quantitative analysis the MS/MS was recombined with MS profile data and peptides that were modified by peptides deamination or oxidation were removed. Peptide features with MOWSE scores <30 (MudPIT scoring) were also excluded.

Progenesis QI (Nonlinear Dynamics, Newcastle, UK) was used to extract mass isotopomer peptide abundance data from MS only spectra. Peak picking was completed on prominent ion features with either 2+ or 3+ charge states. As previously described [17,27], the abundance of the monoisotopic peak (m_0_) and mass isotopomers (m_1_, m_2_, m_3_) were collected covering all the chromatographic peaks for each nonconflicting peptide. Each peptide was matched to the peptides measured in label free quantitation.

### Calculation of Individual Protein Fractional Synthesis Rates

The fractional synthesis rate was calculated in young and aged myoblasts at early and later stages of differentiation by labelling with D_2_O from 0 h - 24 h (early) and 72 h - 96 h (later). Back calculation from peptide mass isotopomer data was performed to quantify the precursor enrichment, as previously described by Hesketh *et al.* [43]. Briefly, the enriched molar fraction of each isotopomer was calculated by subtracting the molar fraction of the unlabelled control peptide from the equivalent D_2_O labelled peptide. The enrichment ratio between m_2_ and m_1_ mass isotopomers was then used to calculate the precursor enrichment.

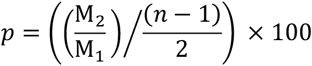

**Equation 1:** *p* = precursor enrichment, M_1_ = enriched molar fraction of m_1_, M_2_ = enriched molar fraction of m_2_, *n* = number of H-D exchange sites.

Deuterium incorporation follows the pattern of an exponential decay. The incorporation of deuterium into newly synthesised proteins causes a decrease in molar fraction of the monoisotopic peak. The rate constant for the decay of the molar fraction of the monoisotopic peak was calculated by a first-order exponential spanning from the start to the end of the 24 h D_2_O labelling period.

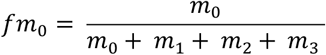

**Equation 2:** *fm*_0_ = molar fraction, *m*_0_ = monoisotopic peak, *m*_1_, *m*_2_, *m*_3_ = mass isotopomers 1 - 3.

The calculation of FSR from the rate constant depends on the number of ^2^H exchangeable H-C bonds. This was calculated by referencing each peptide sequence against standard tables [53]. The FSR of each peptide was derived by dividing the rate constant by the molar percent enrichment of deuterium in the precursor pool and the total number of ^2^H exchangeable H-C bonds in each peptide.

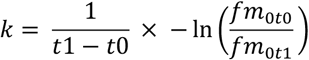

**Equation 3:** *k* = rate constant, *t*0 = first timepoint, *t*1 = end timepoint, *fm*_0*t*0_ = molar fraction at first timepoint, *fm*_0*t*1_ = molar fraction at last timepoint.

The individual protein FSR was reported as the median of each peptide assigned to the specific protein. This value was multiplied by 100 to represent FSR in percent per hour (%/h).

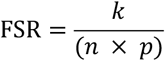

**Equation 4:** *k* = rate constant, *n* = number of H-D exchange sites, *p* = precursor enrichment

### Calculation of Absolute Abundance, Absolute Synthesis Rate and Absolute Degradation Rate

Relative peptide abundance and FSR were converted to absolute values for the total protein in each well. The total protein was extracted and quantified using the BCA™ protein kit, as described above. The total protein (μg/μl) was multiplied by the volume of lysate extracted.

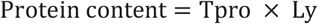

**Equation 5:** Tpro = total protein from BCA assay, Ly = volume of lysate extracted from well.

The normalised relative abundance of proteins was spiked with 50 fmol of yeast alcohol dehydrogenase (ADH1_Yeast) after the tryptic digest. This relative abundance data with ADH1 spike in was multiplied by the total protein content (μg) per well to quantify the absolute protein abundance.

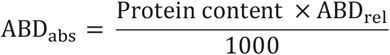

**Equation 6:** ABD_abs_ = absolute protein abundance, ABD_rel_ = normalised relative abundance with ADH1 spike in.

Absolute protein abundance (ABD) was calculated using the protein molecular weight (kDa) specified in the UniProt database and express in absolute values, e.g. μg instead of pmol. We multiplied the absolute protein abundance by the individual protein molecular weight.

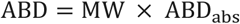

**Equation 7:** ABD = absolute abundance, MW = molecular weight, ABD_abs_ = absolute protein abundance

The absolute synthesis rate was calculated by multiplying the FSR by the absolute protein abundance.

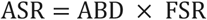

**Equation 8:** ASR = absolute synthesis rate, ABD = absolute abundance, FSR = fractional synthesis rate.

The rate of protein abundance change was calculated by quantifying the difference in absolute protein abundance between the end and beginning of the labelling period. For example, in myoblasts from between 24 h (end) to 0 h (beginning).

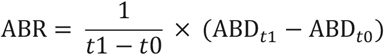

**Equation 9:** ABR = rate of change in absolute protein abundance, ABD = absolute protein abundance.

The absolute degradation rate was calculated by quantifying the difference between the ASR and rate of protein abundance change.

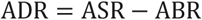

**Equation 10:** ADR = absolute degradation rate, ASR = absolute synthesis rate, ABR = rate of change in absolute protein abundance.

### Creatine Kinase (CK) Assay

On completion of experimentation at 96 h, the cells were washed twice in PBS and lysed with scraping in 250 μl/well of 0.05 M Tris/MES Triton Buffer (TMT; 50 mM Tris MES, 1 % Triton X-100) for 5 min at room temperature. Lysates (20 μl) were added to UV grade 96 well plates (San Jose, CA, USA), followed by 180 μl CK reaction reagent (Catachem Inc., Connecticut, NE). The reaction reagent contained: 30 mmol/l creatine phosphate, 2 mmol/l ADP, 5 mmol/l AMP, 2 mmol/l NAD, 20 mmol/l N-acetyl-L-cystine, 3000 U/l HK (yeast), 2000 U/l G-6-PDH, 10 mmol/l magnesium ions, 20 mmol/l D-glucose, 10 μmol/l Di (adenosine 5’) pentaphosphate, 2 mmol/l ETDA, 0.05 % sodium azide and was buffered at a pH of 6.7. The plate was incubated for 5 min at 37 °C and the change in absorbance was monitored continuously over 20 min using ELISA plate reader (Biotek, USA) at a wavelength of 340 nm.

### Gene Expression Quantification via rRT-PCR

Confluent young and aged C_2_C_12_ myoblasts were washed twice with PBS, lysed with 250 μl/well TRizol reagent for 5 min at room temperature (Sigma Aldrich, Poole, UK). Extracted RNA was transferred to Eppendorf tubes and was isolated by shaking 0.1 ml of chloroform per 0.5 ml of TRizol for 15 s to separate RNA from DNA and protein. Samples were incubated at room temperature for 10 min, centrifuged at 12000 g for a further 15 min at 4 °C. The transparent layer containing chloroform and RNA was extracted and at a ratio of 1:2 isopropanol was added to precipitate the RNA. The samples were incubated for 5 min at room temperature and centrifuged at 12000 g for 10 min at 4 °C. The supernatant was removed, 1 ml of 75 % ethanol was added to the pellet and the samples were centrifuged at 7500 g for 5 min at 4 °C. Following careful decanting, the residual pellet was left to air dry prior to the addition of 20 μl RNA storage solution (Ambion-The RNA Company, Chesire, UK). The RNA was assessed for quality and quantity by adding 1 μl of RNA onto a Nanodrop UV spectrophotometer (Thermo Fisher Scientific, Waltham, USA). Absorbance was measured at 260/280 nm and 260/230 nm to assess RNA purity ratios. For 260/280, a ratio of 2 and for 260/230 of 1.8-2.2 was accepted respectively. RNA concentration was determined, and samples diluted to 70 ng mRNA per gene analysed using a one-step PCR protocol as detailed below.

For RT-PCR, the total reaction volume was 20 μl per sample, containing 11.2 μl master mix (10 μl quantifast SybrGreen, 1 μl primer, 0.2 μl reverse transcriptase) and 8.8 μl RNA (8 ng/μl) sample. The master mix without primers was purchased from Qiagen (Manchester, UK) and primers were synthesised from Primer Design (Southampton, UK). Samples were analysed on rotor-gene Q (Qiagen, Manchester, UK) PCR machine. The amplification protocol was: 1) reverse transcription (10 min at 55 °C), 2) enzyme activation (2 min at 95 °C), 3) denaturation (10 ss at 95 °C) and 4) data collection (60 s at 60 °C) which involved 40 cycles and was followed by melt curve detection. For primer quality control, melt curves were assessed to observe only one peak and uniformity across samples, this was checked with primer design. The amplification efficiencies were analysed, with values 80-100 % proposed as efficient. A threshold was set at 0.1 for all genes to generate C_T_ values which were relativised to 0 h control and reference gene (RP2β) the delta delta C_T_ equation was used [54].

### Cell Fixation and Intracellular Signalling Phosphorylation

To determine early signalling events at the onset of fusion, confluent cells were transferred to DM and harvested at 0, 15, 60, 120 min and 24 h as follows: the cells were washed twice with ice cold PBS prior to addition of 200 μl trypsin for 5 min at 37 °C to detach cells from the gelatin coated plates. The trypsin was neutralised by adding 800 μl GM. The supernatant with cells was centrifuged at 775 g for 5 min at 4 °C. The supernatant was removed leaving a cell pellet which was fixed in 100 μl of 2 % paraformaldehyde and incubated for 60 min at room temperature away from light. Following addition of 500 μl of 100 % methanol, the cells were centrifuged at 500 g for 5 min at 4 °C. The fixatives were removed, the pellet was washed in flow cytometry buffer (PBS + 0.5 % FBS) and centrifuged at 500 g for 5 min at 4 °C. The pellet was resuspended in the flow cytometry buffer and antibodies (Thermo Fisher Scientific, Waltham, USA) were added for: 1) anti-human/mouse phosphor-AKT (S473; APC; 675/25 nm) and 2) anti-human/mouse phosphor-mTOR (S2448; PerCP; 670/LP nm). The cell samples with antibodies were incubated for 60 min at room temperature away from light. The cell samples were washed 3 times in flow cytometry buffer and centrifuged as before. Finally, cells were resuspended in 200 μl flow cytometry buffer and thoroughly mixed before being analysed by flow cytometry on a BD Accuri C6 flow cytometer collecting 2000 events per sample. Compensation of individual fluorescent antibodies in multiple detectors was performed to reduce spectral overlap. Forward scatter and side scatter gating was performed to ensure single populations of cells.

### Morphological Analysis

At 0 h, 24 h, 72 h and 96 h live cell photomicrographs were acquired at 10x magnification using Leica DMI 6000B inverted microscope (Leica Biosystems GmbH, Nussloch, Germany). Myotubes at 72 h and 96 h in the absence and presence of D_2_O were classified as cells that contained ≥ 3 myonuclei/myotube. Myotube number, length, diameter and area was quantified using Image J software (IBIDI, Munich, Germany). Myotube number was counted using the image J cell counter plug in. Myotube length (μm) was measured along the long axis of each tube, diameter (μm) was averaged from measurements at three different (both ends, centre of myotube) equidistant locations and the area (μm^2^) was calculated in Image J from drawing manually around the sarcolemma.

### Statistical Analysis and Graphical Representation

Data are presented as either mean ± standard deviation (SD) or standard error of the mean (SEM) as specifically stated in the figure legends. All statistical analyses were conducted in R (version 4.0.2). Two-way ANOVA were used for comparisons between age (young vs aged) and differentiation period (early vs late). One-way ANOVA was used for comparisons between young alone or aged alone over time. Alternatively, between young and aged at a fixed time-point. For non-proteomic analysis, statistical output was set at P < 0.05. For proteomic comparisons, statistical output was set at P < 0.05 with adjustment based on q-values and FDR of 5 %. Graphical representation was done using either R (version 4.0.2) or Graphpad Prism (version 9.2.0).

## Results

### Experimental Workflow

Deuterium oxide labelling and peptide mass spectrometry were used to investigate the effects of age on the synthesis, abundance, and degradation of individual proteins during the early and late stages of differentiation of murine C_2_C_12_ myoblasts (Figure 1). Consistent with our earlier report [12], this model of replicative ageing was associated with a marked decrease in myotube formation (Figure 2A) and a 6-fold reduction in CK activity (a biochemical marker of differentiation), in aged vs young myotubes (45 ± 2 U.I^−1^ vs 268 ± 18 U.I^−1^: P < 0.001: Figure 2B). Expression of genes involved in myogenesis, including myogenin (200-fold), IGF-I (19-fold) and IGF-II (52-fold; Figure 2C-E) were all also significantly suppressed in aged vs. young (all P < 0.001). Conversely, the mRNA expression of the negative regulator of muscle hypertrophy, myostatin, was 6-fold greater (P = 0.03) in aged compared to young myotubes after 96 h of differentiation (Figure 2F).

**Figure 1.**
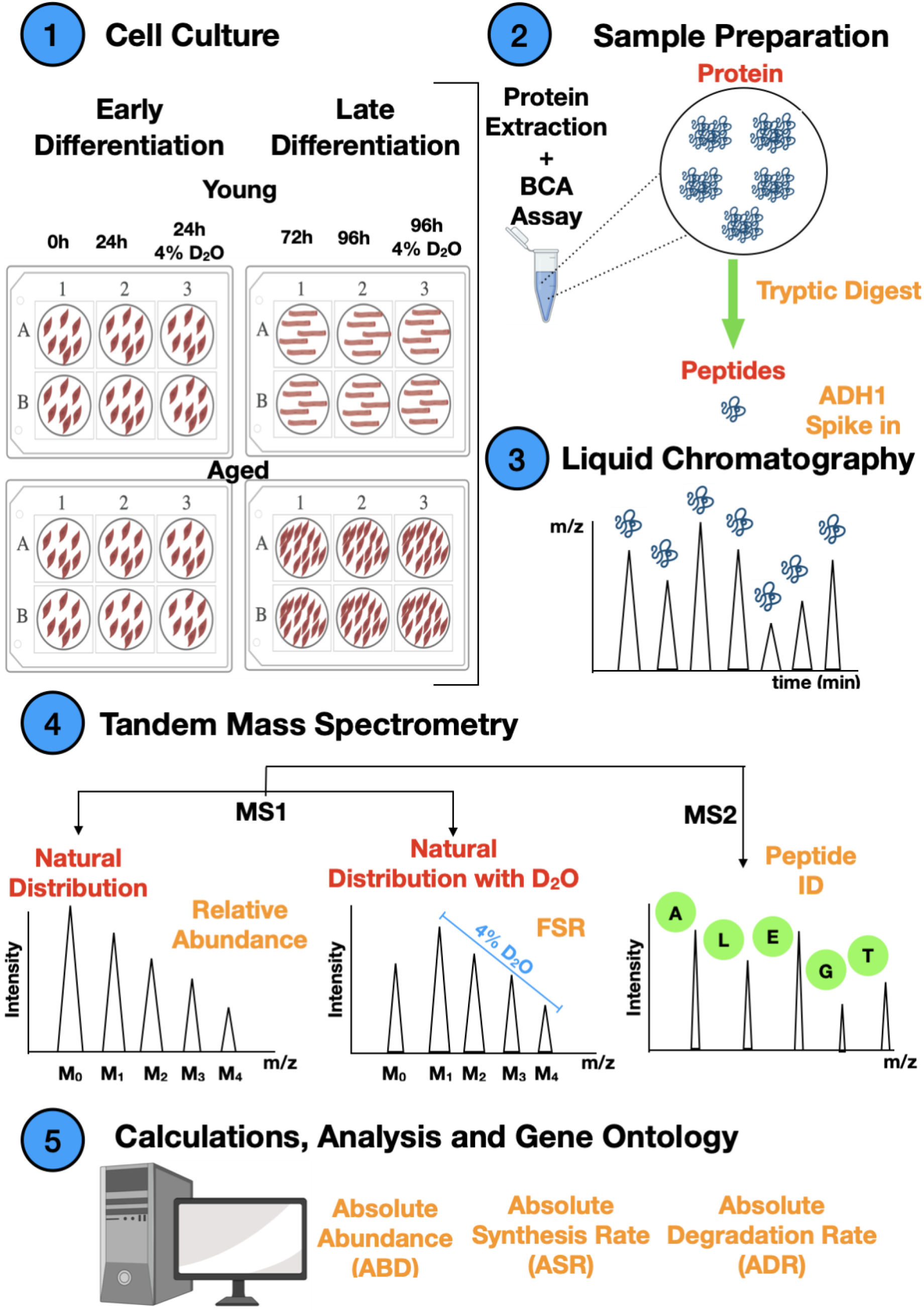
Proteo-ADPT analysis young and replicatively aged myoblast differentiation. Young and aged C_2_C_12_ myoblasts were cultured in the absence or presence of 4 % D_2_O during either early (0-24 h) or later (72-96 h) periods of differentiation. Total protein was extracted and quantified (BCA assay) and proteins were digested into peptides using trypsin. Peptide digests were spiked with 50 fmol of yeast alcohol dehydrogenase-1 (ADH1) and separated by reverse-phase ultra-performance liquid chromatography. Tandem mass spectrometry was used to record MS1 and MS2 mass spectra. MS1 peptide ion mass spectra were used to measure the relative abundance and D_2_O incorporation based on relative distribution of peptide mass isotopomers. The incorporation of deuterium-labelled amino acids in to newly synthesised protein causes a shift in distribution of m_1_, m_2_, m_3_, m_4_… isotopomers, and the m_0_ mass isotopomer declines as a function of protein synthesis. MS2 fragment ion mass spectra of each peptide following were used to infer the amino acid sequence and identify peptides against the UniProt Knowledge database. The Absolute Dynamic Profiling Technique for proteomics (Proteo-ADPT) was used to calculate protein-specific synthesis, abundance and degradation data in absolute (ng) units.

**Figure 2.**
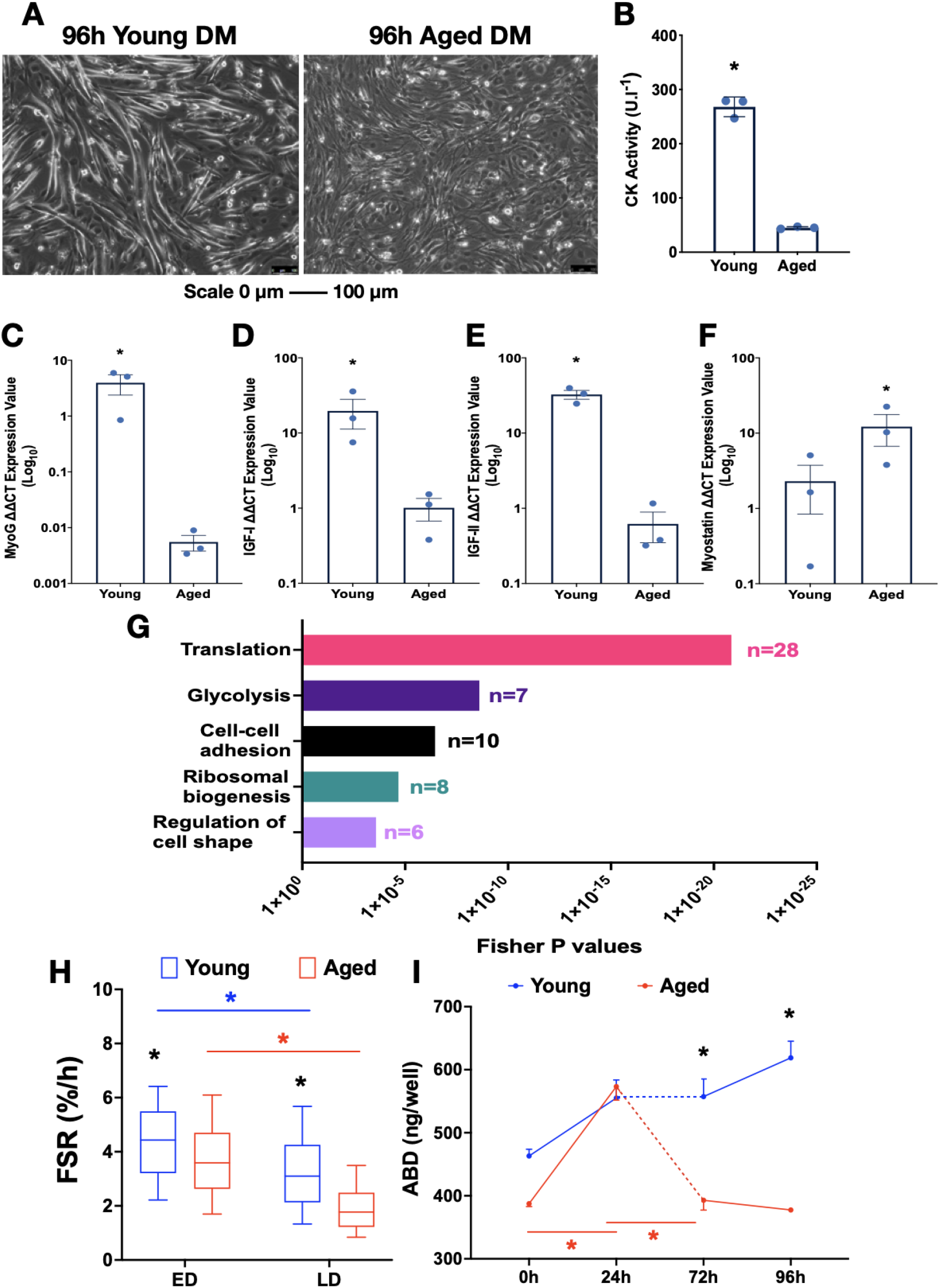
Myotube formation is impaired in replicatively aged C_2_C_12_ myoblasts. **A:** Representative micrographs at 96 h of aged and young myoblast differentiation. Scale 0-100 μm. **B:** Creatine kinase activity in young vs aged at 96 h. **C:** Myogenin gene expression in young vs aged at 96 h. **D:** IGF - I gene expression in young vs aged at 96 h. **E:** IGF - II gene expression in young vs aged at 96 h. **F:** Myostatin gene expression in young vs aged at 96 h. **G:** Gene ontology group for biological processes in 116 proteins matched for abundance, synthesis and degradation. Fisher extract p values quantified using David GO. **H**: Fractional synthesis rates (FSR) of young and aged myoblasts at early differentiation (ED) and late differentiation (LD). **I:** Absolute abundance in young vs aged myoblasts at 0 h, 24 h, 72 h, 96 h. Data presented as mean ± SD for creatine kinase and SEM for gene expression and abundance. FSR data presented as median and interquartile range. In the gene expression data, the aged was relativised to young basal Ct values, the reference gene RP2β and the 0 h timepoint. Significance (*) between young and aged was set at P < 0.05. All experiments N=3 in duplicate.

Consistent with our previous findings [21], supplementation of media with D_2_O had no observable effect on myoblast viability (Figure S1A) or myotube (Figure S1B) formation; that is, the length, diameter and area of myotubes were equivalent when grown in the absence or presence of D_2_O in young cells (mean ± SEM; length, diameter and area: 65 ± 2 μm, 6.4 ± 0.1 μm and 414 ± 14 μm^2^; Figure S1C). Since replicative ageing results in the prevention of myotube formation [12], myotube morphology could not be assessed, however all cells remained viable in the absence or presence of D_2_O (Figure S1A/B).

Proteins were extracted from control and deuterium-labelled cell cultures at 0 h, 24 h, 72 h and 96 h after transfer to differentiation media and submitted to proteomic analysis. After stringent filtering to remove proteins that were not detected in all (n=24) samples, the analysis included 611 proteotypic peptides belonging to a total of 116 individual proteins (Supplementary File 1). The top-ranked biological processes (Fisher P < 0.001; Figure 2G) amongst the proteins studied, included: translation (n=28), glycolysis (n=7), cell-cell adhesion (n=10), ribosomal biogenesis (n=8) and regulation of cell shape (n=6). Protein fractional synthesis rates (FSR) were calculated from time-dependent changes in peptide mass isotopomer distribution that reflect the incorporation of deuterium into newly synthesised proteins [55] during either early (0 h - 24 h) or later (72 h - 96 h) periods of differentiation. Irrespective of age, mixed protein FSR was significantly greater during early compared to late differentiation, which is consistent with data reported in low-passage myoblasts and myotubes [21]. However, the decline in mixed protein FSR between early and late period of differentiation was significantly greater in aged cells (P < 0.05 interaction between differentiation period and cell age; Figure 2H). The median and interquartile range of protein FSR was significantly (both P < 0.05) greater in young myoblasts (4.4 ± 2.3 %/h) and in young myotubes (3.1 ± 2.1 %/h) compared to aged myoblasts (3.6 ± 2.0 %/h) and aged myotubes (1.7 ± 1.3 %/h). To facilitate analysis using the absolute dynamic profiling technique for proteomics (Proteo-ADPT) reported previously [52], relative protein abundances and FSR data were normalised against a spiked standard (50 fmol yeast alcohol dehydrogenase-1) and individual protein data were multiplied by the total protein content (Table S3) of the respective well, to report protein abundances and synthesis rates in absolute (ng/well and ng/well/h) units.

The total protein abundance in each well tended (P = 0.09) to exhibit an interaction between differentiation stage and replicative age, and there were statistically significant (P < 0.05) main effects for both differentiation period and replicative age. In young cells, differentiation was associated with a consistent rise in protein content from 463 ± 10 ng/well at 0 h through to 618 ± 65 ng/well (~25 % increase; P = 0.052) after 96 h of differentiation. In contrast, aged cells exhibited a robust 32 % increase (from 388 ± 40 ng/well to 573 ± 63 ng/well: P = 0.007) in protein content during early differentiation (0 h - 24 h) but the protein content of aged cells then declined back to baseline levels during the later period of differentiation (392 ± 43 ng/well at 72 h and 377 ± 38 ng/well at 96 h; Figure 2I).

### Protein-specific responses in young and aged myoblasts during early and late differentiation

Two-way analysis of variance was used to investigate the interaction between replicative ageing and early vs late periods of myoblast differentiation in the rates of abundance change (ABR; ng/well/h) of each protein (Figure 3). In all cases, a threshold of P < 0.05 and a false discovery rate of 5 % was used to infer statistical significance.

**Figure 3.**
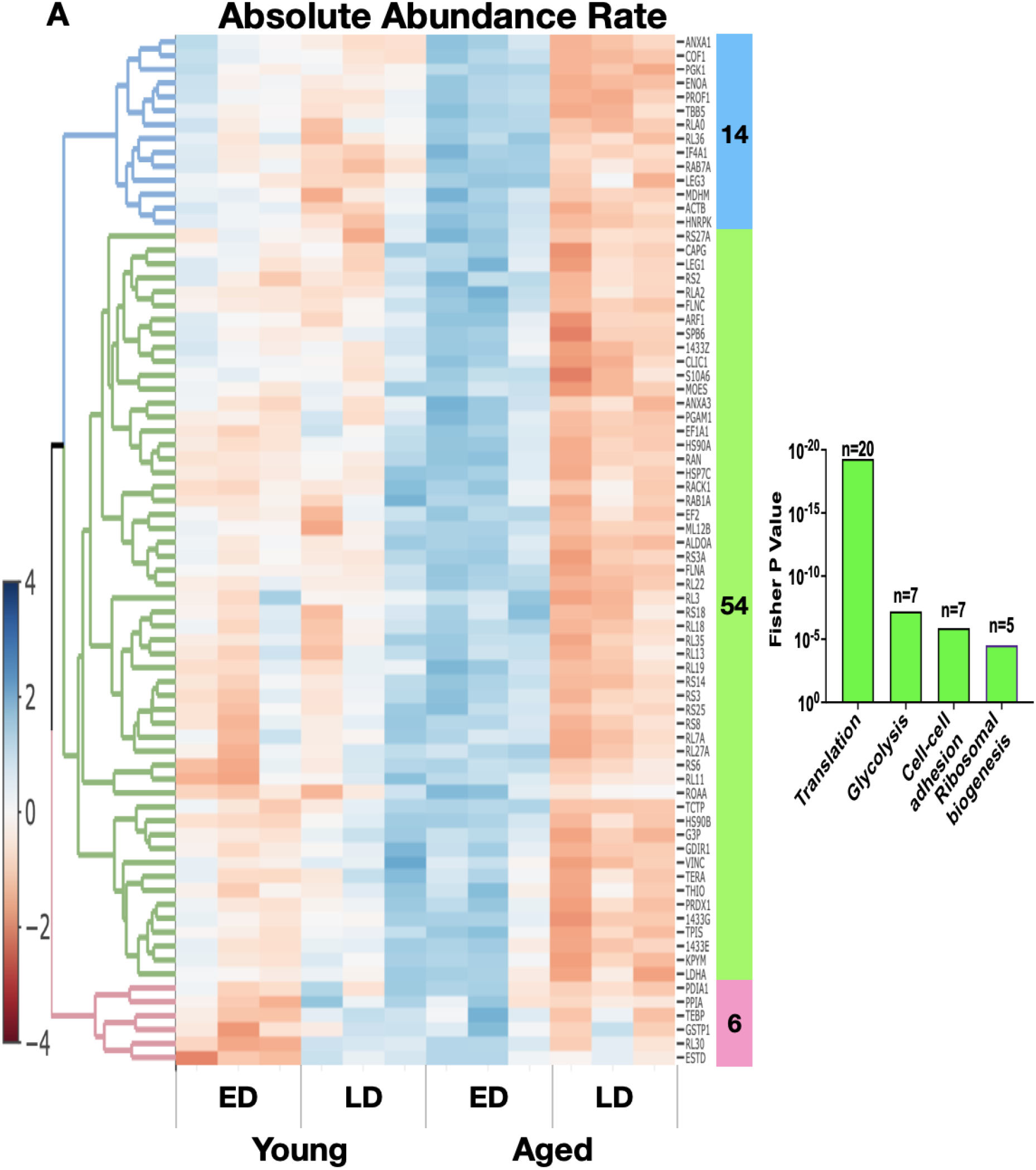
Protein rate of abundance change in young and aged cells during early and later differentiation. Heatmap of proteins with significant interaction (P < 0.05: FDR 5 %) between age and early/later differentiation (ED/LD) absolute abundance rates (ABR) with 74 proteins. Blue increased and red decreased expression. Proteins were clustered based on dendrogram on left of heatmap and to the right of the heatmap, top-ranked biological processes of 54 proteins clustered in green displayed in bar graph. All experiments N=3 in duplicate.

The majority (63 %; 74/116) of proteins analysed exhibited significant interactions (age x stage of differentiation) in ABR. In young cells protein accretion was consistent and the majority of individual proteins exhibited positive ABR values that were similar during early and late periods of differentiation. In aged cells 54 proteins exhibited positive ABR values that were greater than those of young myoblasts during early differentiation. In most cases the abundance of these proteins had been significantly less in aged cells at the 0 h timepoint (Figure 3A), therefore, the greater ABR compensated for the lower starting abundance and brought the proteome profile of aged cells more closely in line with that of young myoblasts at the end of the early differentiation period. The protein responses specific to aged cells encompassed several significantly (all Fisher P < 0.001) enriched biological processes, including translation (n=20), glycolysis (n=7), cell-cell adhesion (n=7) and ribosomal biogenesis (n=5). In aged myoblasts several proteins associated with translation initiation and elongation (eukaryotic initiation factor 4A-I, IF4A1; elongation factor 1-alpha 1, EF1A1; and elongation factor 2, EF2) exhibited significantly greater ABR than young cells, which rescued the abundance to the level of the young. In addition, 18 ribosomal proteins (10 large ribosomal subunits and 8 small ribosomal subunits) exhibited greater gains in abundance during early compared to late differentiation in aged cells. Three chaperone proteins (heat shock protein 90 alpha; HS90A, heat shock protein 90 beta, HS90B, heat shock cognate 71 kDa protein; HSP7C) also displayed greater gains in abundance during early compared to late differentiation in aged cells and similar significant patterns of change were evident in the abundance of enzymes involved in glycolytic energy metabolism and cell adhesion.

### Proteo-ADPT analysis of early differentiation

The differences in ABR between young and aged cells (Figure 3) were further interrogated using the absolute dynamic profiling technique for proteomics (Proteo-ADPT). Two-way ANOVA of the absolute abundance (ABD), synthesis rate (ASR) and degradation rate (ADR) was used to investigate differences between young and aged cells from the onset (0 h) to the end (24 h) of the early differentiation process (Figure 4).

**Figure 4.**
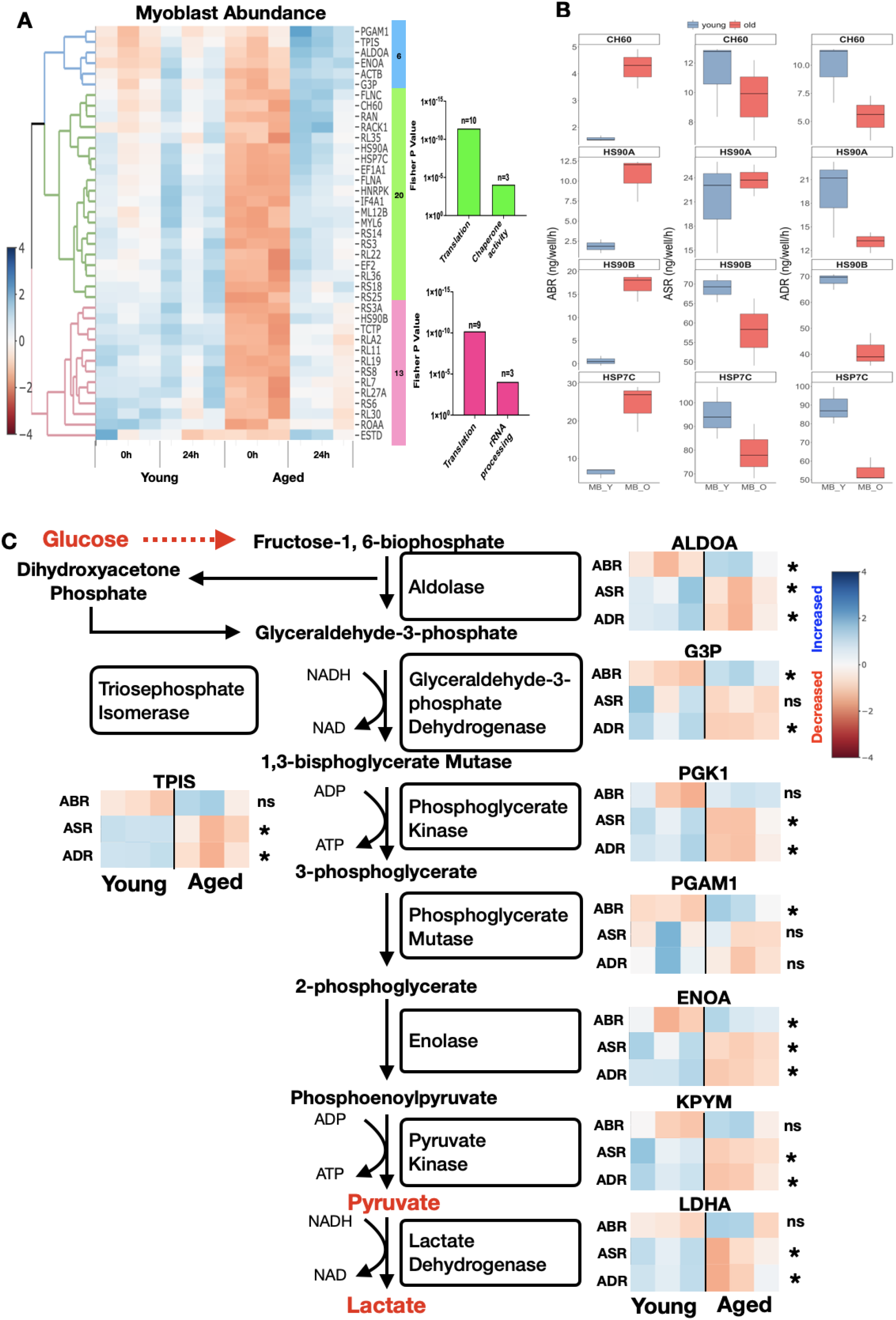
Proteo-ADPT in young and replicatively aged cells during early differentiation. **A:** Heatmap of proteins with significant interaction (P < 0.05: FDR 5 %) between age and cell absolute abundance (ABD) with 49 proteins. Blue increased and red decreased expression. Proteins were clustered based on dendrogram on left of heatmap and to the right of the heatmap, top-ranked biological processes were displayed in bar graphs with corresponding number of proteins. **B**: Selected individual protein rate of abundance change (ABR), absolute synthesis (ASR) and degradation rates (ADR) in young and aged myoblasts. **C**: Proteo-ADPT model (ABR, ASR, ADR) on enzymes identified in glycolysis pathway. Blue increased and red decreased expression. Significance between young and aged indicated with *. All experiments N=3 in duplicate.

Thirty-nine proteins exhibited significant (P < 0.05: FDR 5 %) interactions in ABD between age and time (Figure 4A), and the biological processes enriched (Fisher P < 0.001) amongst these proteins included chaperone activity (n=4), glycolysis (n=5) and translation (n=19). At the beginning (0 h) of the labelling period the chaperone proteins HS90A, HS90B, HSP7C and 60 kDa heat shock protein, mitochondrial (CH60) were significantly more abundant in young compared to aged cells but this difference was no longer evident at the end (24 h) of the labelling period (Figure 4A). That is, coinciding with the greater rate of abundance change (Figure 3), the aged cells recovered the abundance of these proteins to the level measured in young cells by the 24 h time point. Interestingly, the synthesis rates of these chaperone proteins were similar in young and aged cells and the greater rate of increase in abundance was due to significantly lesser degradation of the chaperone proteins in aged cells. For example, the abundance of HS90A increased at a rate of 10.62 ± 1.60 ng/well/h in aged which was significantly greater than 1.86 ± 0.49 ng/well/h in young cells (Figure 4B). The ASR in young (21.23 ± 3.40 ng/well/h) vs aged (23.69 ± 1.13 ng/well/h) were equivalent and the ADR was significantly (P = 0.05) greater in young vs aged cells (19.37 ± 2.93 ng/well/h vs 13.06 ± 0.76 ng/well/h).

The abundance of several glycolytic enzymes was also less in aged than young cells at the beginning (0 h) of the labelling period and, similar to the chaperone proteins, these differences in abundance were recovered in aged cells by the 24 h time point. Unlike chaperone proteins, however, age-related differences in abundance changes of glycolytic enzymes were not entirely explained by differences in protein degradation rates. Eight glycolytic enzymes exhibited significant differences between young and aged cells in either abundance (Figure 4A), synthesis and/or degradation rates (Figure 4C). For example, the greater rate of increase in fructose-bisphosphate aldolase A (ALDOA) and alpha-enolase (ENOA) in aged cells was driven by greater synthesis rates (Figure 4C: Table S4). Conversely, the greater gain in abundance of glyceraldehyde-3-phosphate dehydrogenase (G3P) in aged cells were due to lower degradation rates of G3P in aged vs young cells. Phosphoglycerate mutase 1 (PGAM1) abundance response was due to non-significant differences in synthesis and degradation in aged compared to young (Figure 4C: Table S4). Lastly, there was no difference in ABR between young and aged for the proteins triosephosphate isomerase (TPIS), phosphoglycerate kinase 1 (PGK1), pyruvate kinase (KPYM) and L-lactate dehydrogenase A chain (LDHA), but the turnover of these proteins was significantly greater in young myoblasts (Figure 4C: Table S4).

The 19 proteins associated with translation processes, included 9 proteins of the large ribosomal subunit, 7 proteins of the small ribosomal subunit and 3 proteins that regulate either the initiation or elongation phase of translation (Figure 5A). At the onset of differentiation, young myoblasts had a significantly greater abundance of ribosomal proteins, and the abundance of ribosomal proteins remained stable throughout the early 0 h - 24 h differentiation period. In aged cells the abundance of ribosomal proteins increased during early differentiation so that by 24 h, only 2 proteins (60S ribosomal protein L11; RL11 and 60S acidic ribosomal protein P2; RLA2) of the aforementioned 19 remained significantly more abundant in young myoblasts. This dramatic change in aged myoblasts from 0 h to 24 h was underpinned by significant positive gains in the abundance in each of the 19 proteins (Figure 5A: Table S5). The majority (n=13: Figure 5A) of changes in protein abundance were not explained by heightened synthesis in aged cells. Indeed, EF1A1, RL11, 60S ribosomal protein L19 (RL19), 60S ribosomal protein L22 (RL22) and 40S ribosomal protein S25 (RS25) exhibited greater ASR in young cells but the gain in the abundance of these proteins was greater in aged cells. Accordingly, the degradation rates of RL11, 40S ribosomal protein S6 (RS6), 40S ribosomal protein S8 (RS8), RL19, RL22, EF2, EF1A1, IF4A1 and RS25 were significantly less in aged cells and accounted for the greater gains in the abundance of these proteins compared to young myoblasts. Only RL11 had significantly greater abundance and turnover in young cells, whereas 40S ribosomal protein S3A (RS3A) had significantly greater turnover in aged cells. Five proteins (EF1A1, RL11, RL19, RL22, RS25) had significantly greater turnover in young compared to aged cells and did not exhibit differences in protein abundance (i.e. abundance was at equilibrium between 0 h and 24 h of differentiation), which may suggest the quality of these proteins was diminished in aged cells.

**Figure 5.**
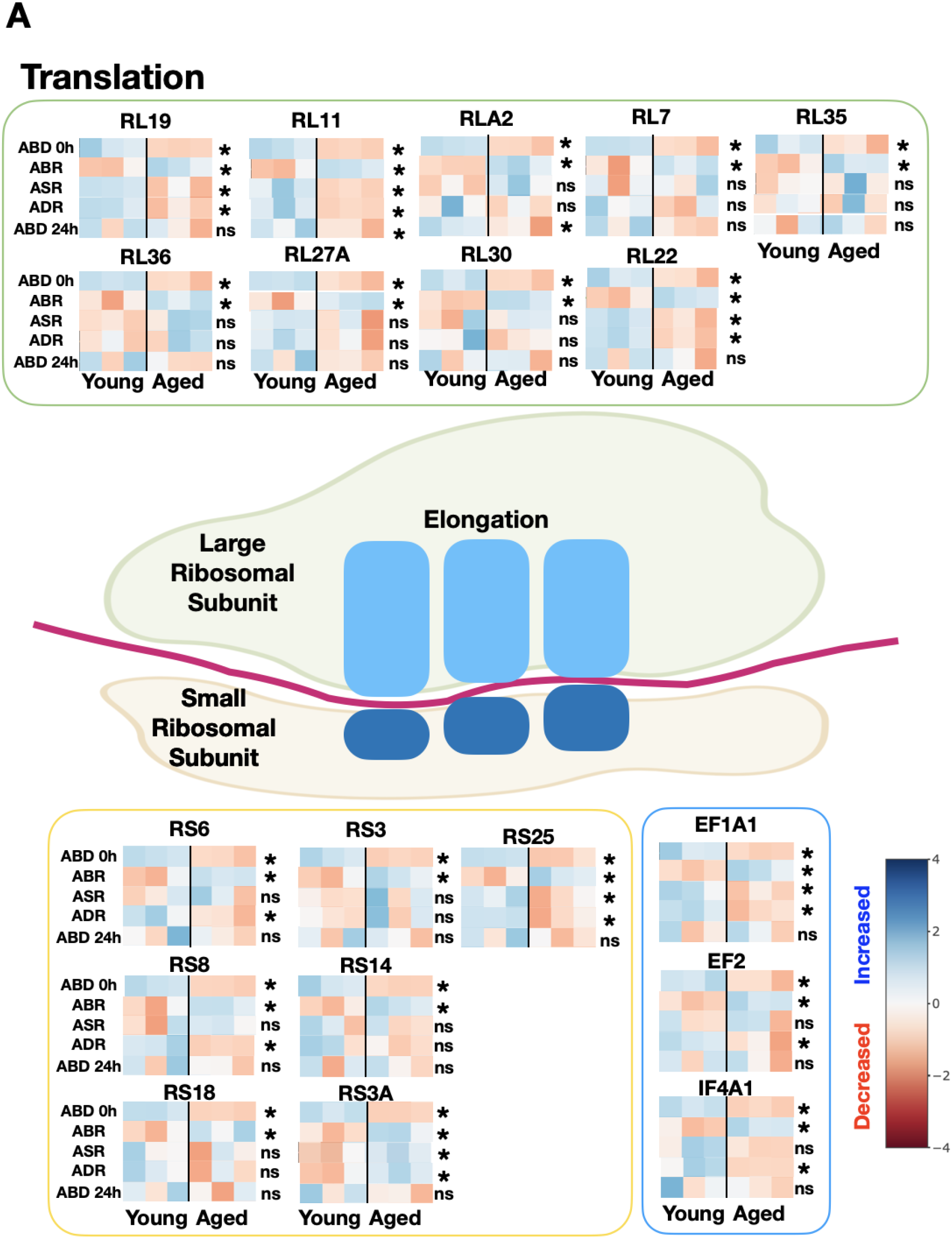
Proteo-ADPT of translation proteins between young and replicatively aged cells during early differentiation. Absolute abundance and Proteo-ADPT model (ABR, ASR, ADR) on ribosomal proteins identified that regulate translation. Blue increased and red decreased expression. Significance between young and aged indicated with *. All experiments N=3 in duplicate.

### Proteo-ADPT analysis of late differentiation

During the later period (72 h – 96 h) of differentiation, the abundance of 6 proteins KYPM, HS90A, HS90B, filamin-A (FLNA), EF1A1 and 6-phosphogluconate dehydrogenase (6PGD) exhibited a statistical interaction (P < 0.05: FDR 5 %) between time and age (Figure 6A). Generally, these proteins were significantly less abundant in aged compared to young cells at the 72 h time point and further reduced in abundance specifically in aged cells during the 72 h to 96 h period.

**Figure 6.**
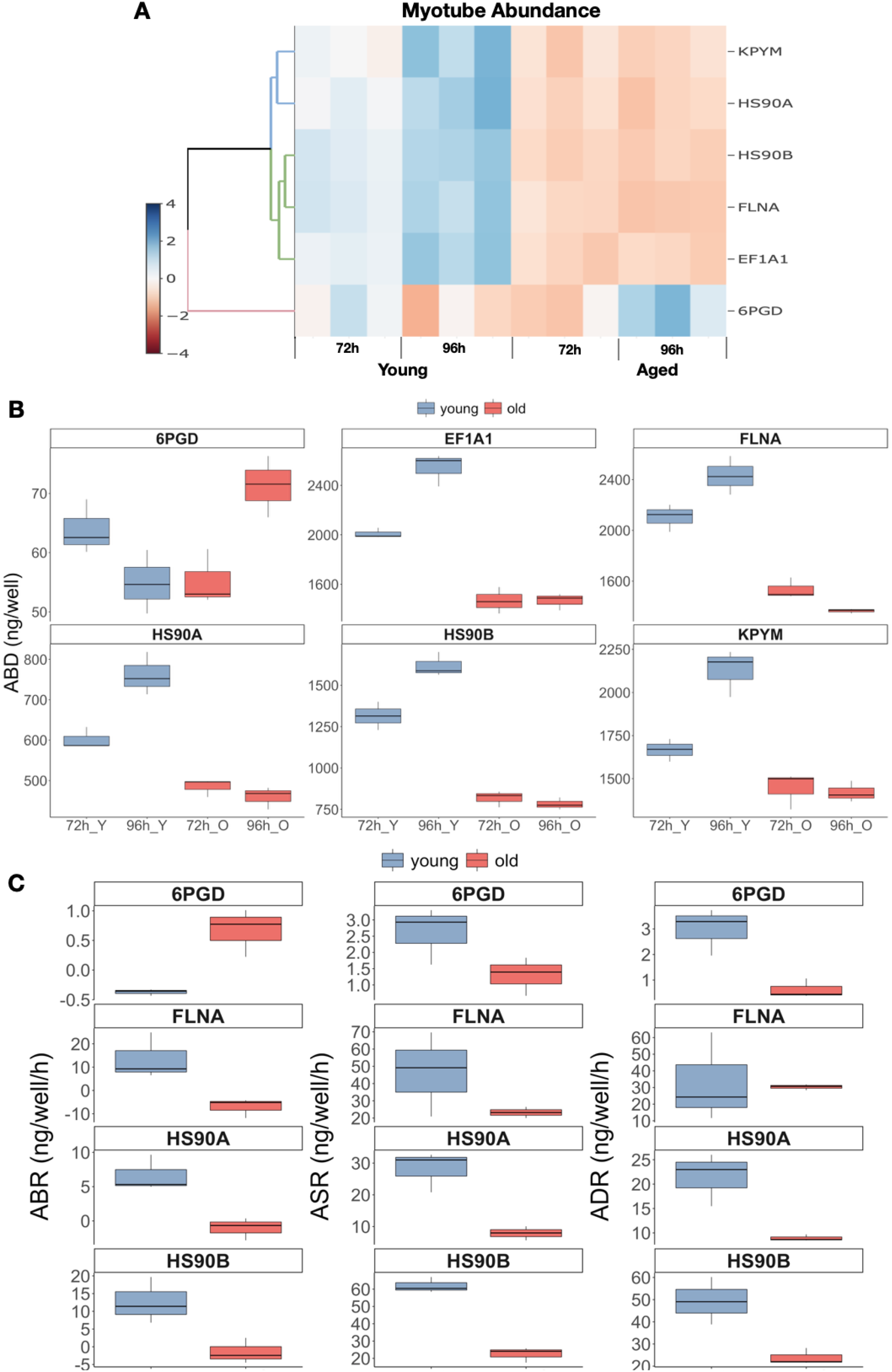
Proteo-ADPT in young and replicatively aged cells throughout later differentiation. **A:** Heatmap of proteins with significant interaction (P < 0.05: FDR 5 %) between age and cell absolute abundance (ABD) with 6 proteins. Blue increased and red decreased expression. Proteins were clustered based on dendrogram on left of heatmap. **B**: Individual proteins ABD in young and aged myoblasts at late differentiation. **C**: Selected individual protein rate of abundance change, absolute synthesis and degradation rates in young and aged myoblasts at late differentiation. All experiments N=3 in duplicate.

In young cells the abundance of HS90A increased (positive ABR of 6.65 ± 1.51 ng/well/h; Figure 6B) underpinned by a positive net balance between synthesis and degradation (ASR = 28.14 ± 3.70 ng/well/h vs ADR = 21.49 ± 3.13 ng/well/h). Conversely, in aged cells the degradation rate of HS90A (8.94 ± 0.38 ng/well/h) outweighed the rate of synthesis (7.89 ± 1.28 ng/well/h) meaning that the turnover of HS90A was also significantly less compared to young cells. Similarly, HS90B abundance increased at a rate of 12.63 ± 3.76 ng/well/h, in young cells which was significantly different (P < 0.05) from the rate of loss (−1.47 ± 2.07 ng/well/h) in abundance of HS90B in aged cells (Figure 6C). In young cells HS90B was synthesised at a rate of 61.98 ± 2.57 ng/well/h and degraded at a rate of 49.35 ± 6.17 ng/well/h. In aged cells, HS90B rates of synthesis (22.46 ± 2.51 ng/well/h) and degradation (23.93 ± 2.13 ng/well/h) were lower compared to young cells and balanced in favour of a reduction in HS90B protein abundance. FLNA abundance gain was 20.89 ± 7.84 ng/well/h in young whereas, in aged the abundance rate decreased at −7.03 ± 2.39 ng/well/h (Figure 6C). Similar to the heat shock proteins, HS90A and HS90B, the contrasting abundance change in FLNA was a result of greater synthesis than degradation in young (ASR = 133.87 ± 17.68 vs ADR = 112.98 ± 25.20 ng/well/h) compared to greater degradation than synthesis in aged (ASR = 30.70 ± 3.51 ng/well/h vs ADR = 31.08 ± 3.46 ng/well/h). In contrast to the aforementioned proteins, the abundance of 6PGD decreased in young and increased in aged cells during the late differentiation period (Figure 6B). 6PGD displayed a negative rate of abundance change of −0.37 ± 0.03 ng/well/h in young and a positive rate of change 0.67 ± 0.23 ng/well/h in aged (Figure 6C). The decline in abundance was driven by greater degradation than synthesis in young (ASR = 2.62 ± 0.51 ng/well/h vs ADR = 2.99 ± 0.53 ng/well/h), whereas synthesis was greater than degradation in aged cells (ASR = 1.30 ± 0.34 ng/well/h vs ADR = 0.63 ± 0.22 ng/well/h).

### Gene expression and signalling during early differentiation

IGF-I and myostatin gene expression were measured at 0 h and 24 h time points, and the phosphorylation of Akt (serine 473) and mTOR (serine 2448) were measured at 0 h, 15 min, 60 min, 120 min, and 24 h after cells were transferred into differentiation media. Myostatin gene expression was significantly (P < 0.05, main effect of age; Figure 7A) greater in aged compared to young at 0 h (34-fold) and 24 h (12-fold). Conversely, IGF-I gene expression was 9-fold greater (P < 0.05, interaction effect; Figure 7B) in young compared to aged cells at the 0 h timepoint. IGF-I gene expression significantly (both P < 0.05) increased by 7-fold over the 24 h period in both young and aged cells but IGF-I expression was consistently 10-fold greater (P < 0.05) greater in young vs aged. There were also significant interactions (P < 0.05) between time and age for both Akt and mTOR phosphorylation (Figure 7B and D). Coinciding with the changes in IGF-I gene expression, young cells exhibited significantly (P < 0.05) greater phosphorylation of signalling intermediates compared to aged cells at both 0 h (Akt = 2.7-fold: mTOR = 2.0-fold) and 24 h (Akt = 9.9-fold: mTOR = 10.9-fold). However, at the 15 - 120 min time points, there was no statistically significant differences between young and aged cells in either Akt or mTOR phosphorylation.

**Figure 7.**
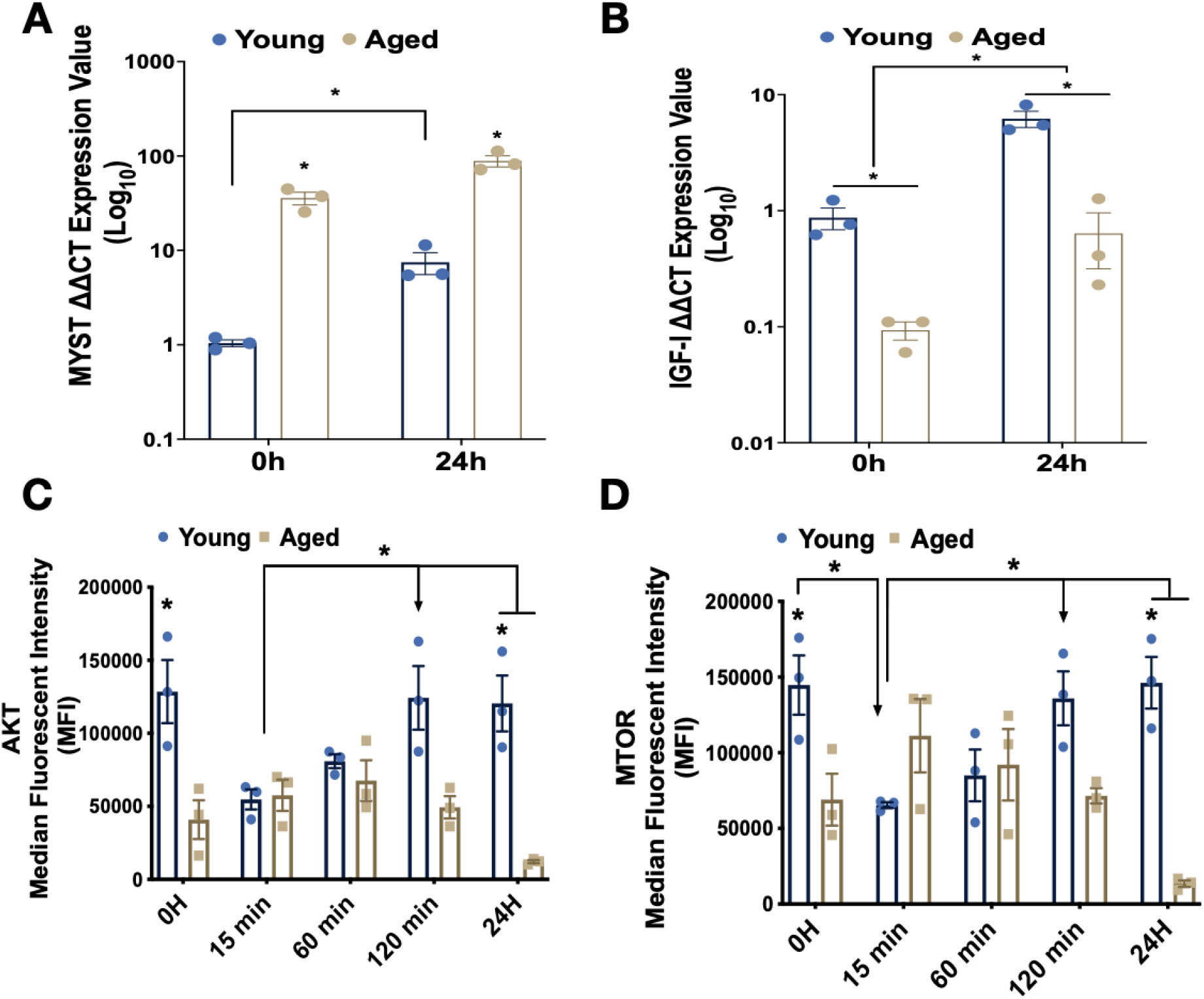
Myostatin expression and IGF-I – Akt – mTOR signalling in young and replicatively aged cells during early differentiation. **A:** IGF - I gene expression in young vs aged at 0 and 24 h. **B**: Akt phosphorylation in young vs aged myoblasts at 0 h, 15 min, 60 min, 120 min and 24 h **C:** mTOR phosphorylation in young vs aged myoblasts at 0 h, 15 min, 60 min, 120 min and 24 h. Data presented as mean ± SEM for **A-C**. All experiments N=3 in duplicate.

## Discussion

Ageing muscle undergoes a loss of mass and strength, phenotypes which partially rely on the ability of muscle stem cells to regenerate the muscle. We used Proteo-ADPT [52] to investigate the absolute abundance, synthesis, and degradation rates of individual proteins in low passage ‘young’ and replicatively-aged myoblasts during early and late periods of differentiation. Young cells exhibited a steady pattern of growth, protein accretion and fusion throughout the first 96 h of differentiation. In contrast, aged myoblasts exhibited markedly different proteome responses during early differentiation and failed to gain protein mass or undergo fusion during later differentiation (Figure 8). Our novel Proteo-ADPT data provide new evidence suggesting that the maturation of the proteome may be retarded in aged myoblasts at the onset of differentiation but quickly catches up with the young cells during the first 24 h period. However, this ‘catch up’ process in aged cells is not accomplished by higher levels of protein synthesis. Instead, a lower level of protein degradation in aged cells was responsible for the gains in protein abundance. As differentiation progressed, aged cells also did not fuse in to myotubes and their protein content declined. Our novel data point to a loss of proteome quality as a precursor to the lack of fusion of aged myoblasts and highlights dysregulation of protein degradation, particularly of ribosomal and chaperone proteins, as a key mechanism that may contribute to age-related declines in the capacity of myoblasts to undergo differentiation.

**Figure 8.**
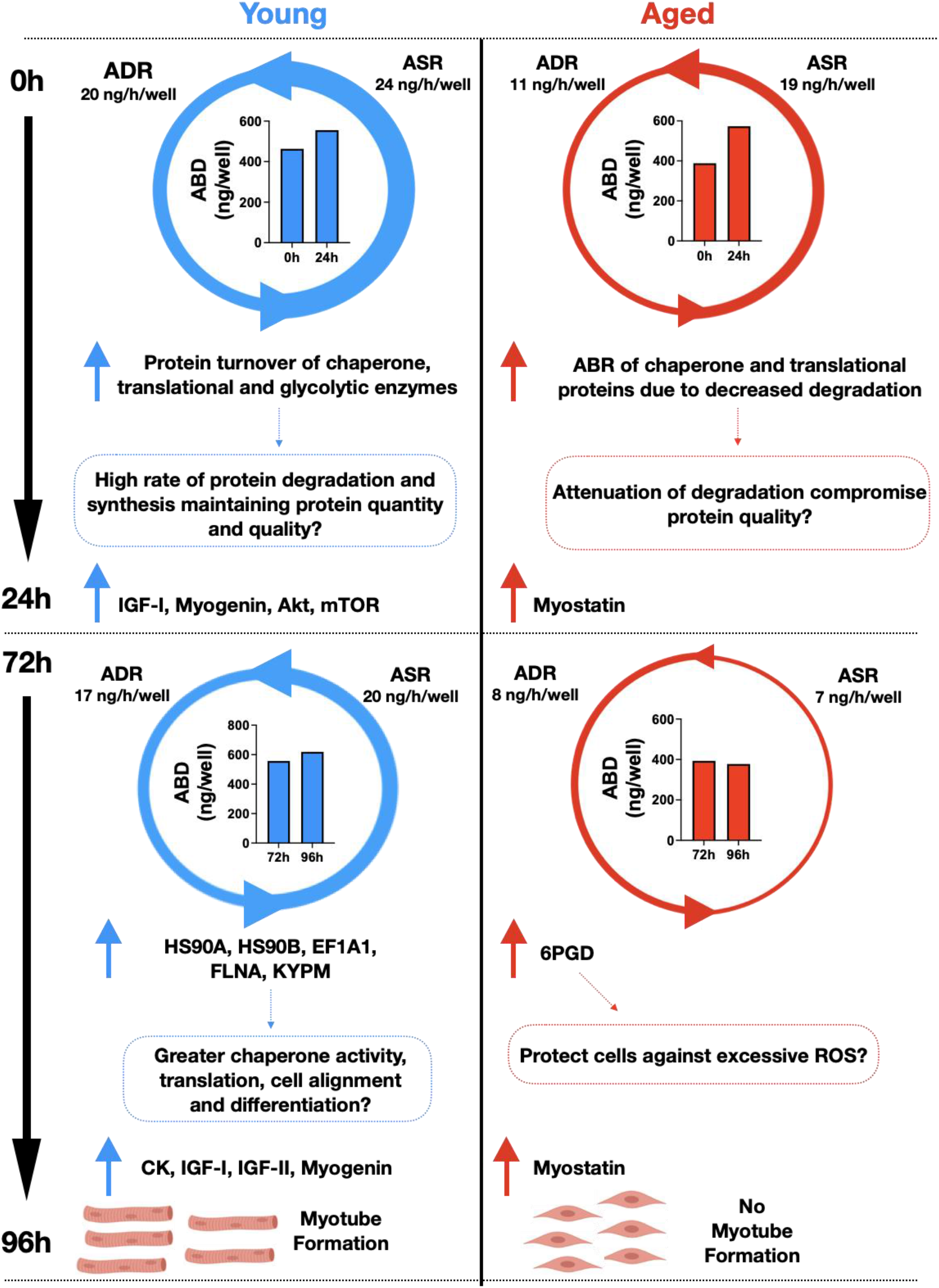
Summary of young compared to replicatively aged cells during early and late myoblast differentiation. The summary of our findings on young myoblasts are on the left column and aged on the right column. Downwards arrows covers early and late myoblast differentiation. Size of circular arrows corresponds to degradation and synthesis rates. ABD; absolute abundance, ADR: absolute degradation rates, ASR: absolute synthesis rates, ABR: absolute rates of abundance change, IGF-I: insulin-like growth factor I, IGF-II: insulin-like growth factor II, Akt: protein kinase B, mTOR: mechanistic target of rapamycin, HS90A: heat shock protein 90 alpha, HS90B: heat shock protein 90 beta, EF1A1: elongation factor 1-alpha 1, FLNA: filamin-A, KYPM: pyruvate kinase, 6PGD: 6-phosphogluconate dehydrogenase, ROS: reactive oxygen species, CK: creatine kinase.

Ubaida-Mohien *et al*. [32], report decreases in abundance of ribosomal proteins and elevated abundance of proteins involved in proteostasis as a function of advancing age in human skeletal muscle. Our data suggest similar changes may extend to the proteome of muscle precursor cells and underpin age-related declines in regenerative capacity and we add insight regarding the contribution of synthesis and degradation to an age-related differences in protein abundance. Studies in model organisms [34,35], experimental models [38,39] and humans [40,41] consistently report protein turnover declines as a function of advancing age and we also report a general drop in mixed-protein synthesis in replicatively aged myoblasts. Modelling of data from studies of ageing in C. elegans [45] predicts that advancing age leads to a collapse in proteostasis, which can be rescued by overexpression of chaperones or protein degradative systems. However, the gradual age-related decline in protein turnover in *C. elegans* is not generalised to the whole proteome and rather is associated with greater heterogeneity of turnover rates amongst proteins [35]. Herein ribosomal subunits and chaperone proteins exhibited significantly altered abundance and synthesis throughout the differentiation of replicatively aged myoblasts. Coinciding with our data, identical ribosomal and chaperone proteins also exhibited reduced abundance throughout the *C. elegans* lifespan and the turnover rates were also heterogeneous [35]. We report declines in both synthesis and degradation in aged cells at the later stages of differentiation and aberrant changes in the abundance of ribosomal and chaperone proteins in aged myoblasts in early differentiation. Proteo-ADPT enabled us to distinguish differences in the contribution of synthesis and degradation to changes in protein abundance and turnover rate in young and aged cells and the ribosomal and chaperone abundance alterations in aged were underpinned by differences in protein degradation rather than synthesis. Ribosomal proteins are degraded by the UPS system [56] and heightened activation of the 20S proteasome is able to extend lifespan in *C. elegans* [50]. Indeed, age-related declines in proteasome abundance and activity precedes the loss of ribosomal subunit stoichiometry in the brain [57]. Together these findings suggest an age-related reduction in proteasome activity precedes the loss in ribosome stoichiometry and is a precursor to the widely overserved general decline in protein turnover.

Chaperone proteins are integral to proteostasis and regulate myoblast differentiation by interacting with key cell signalling pathways [58]. Substrate proteins, or clients of the alpha (HS90A) and beta (HS90B) isoforms of heat shock protein 90 (HSP90) include MRFs, such as MyoD and myogenin [47], and Akt, which are all required for myogenic differentiation [59,60]. We found that aged myoblasts had a significantly lower abundance of HS90A and HS90B at the onset of differentiation, and although the abundance of HS90A and HS90B recovered to levels equivalent to young cells after 24 h of differentiation, this accelerated gain in abundance was not achieved by a rise in the synthesis rate of HS90A or HS90B. Instead, a lack of degradation of HSP90 isoforms in aged cells was associated with the accumulation in protein abundance during early differentiation, which may suggest the quality of these chaperone proteins in aged cells was suboptimal at the 24 h time point. During later differentiation, the abundance of HS90A and HS90B was reduced in aged cells that failed to undergo fusion and form myotubes, unlike in young cells, which exhibited a significant gain in abundance and formed myotubes. Pharmacological inhibition of HSP90 by Geldanamycin treatment does not alter the abundance of HSP90 but disables the ability of HSP90 to release client proteins, which causes client proteins to be degraded [61]. Inhibition of HSP90 retards muscle regeneration *in vivo* and stalls protein accretion and the fusion of myoblasts in to myotubes *in vitro* [59] in a manner that closely resembles our findings in replicatively aged myoblasts. Akt is an HSP90-dependent kinase and the interaction between Akt and HSP90 modulates the phosphorylation status of Akt [60]. The phosphorylation of Akt rose steadily during early differentiation in young myoblasts but this time-dependent response failed in replicatively aged cells (Figure 7C). Akt function is required for myoblast differentiation [59] and our findings suggest disruption to the turnover of HS90A and HS90B may be particularly associated with diminished Akt phosphorylation and the failure to form myotubes in aged cells. FLNA acts as a mechanotransducer during cell migration and differentiation by binding to actin, focal adhesions and sarcomeric Z-disc [62,63]. During cell alignment, which precedes fusion, filamins rapidly unfold and dictate movement at the leading cell edge [63] and this unfolding is driven by binding to HSP90 [64]. Therefore, this age-related reduction in FLNA and HSP90 abundance could be associated with the lack of fusion during later differentiation.

Heat shock protein 70 (HSP70) and CH60 work cooperatively with HSP90 [61] and we report HSP7C and CH60 exhibit age-dependent differences in protein abundance, synthesis and degradation similar to the patterns exhibited by HS90A and HS90B (Figure 4B). Diminished expression of either the inducible 70 kDa heat shock protein (HSPA1) or the closely related HSP7C disrupts differentiation of C_2_C_12_ myoblasts [65] similar to the effects HSP90 inhibition [47], however the interacting proteins and kinase pathways involved are likely to be different. While HSP90 impacts Akt-related signalling, the MAPK p38α is the key client of the HSP70 proteins [65]. P38 MAPK interacts with the transcription factors MEF2 and MyoD to regulate myoblast differentiation [29] and HSP70 is required for myoblast differentiation *in vitro* and muscle regeneration *in vivo* [65]. Loss of HSP70 can be rescued by either enhancing p38 MAPK or through proteasome inhibition [65], which implies that HSP70 stabilises and prevents the degradation of p38 MAPK. The key role of HSP70 in myoblast differentiation is consistent with the improvement in muscle regeneration after damage caused by eccentric contractions when HSP70 is overexpressed compared to aged wild-type mice [66]. CH60 is a key mitochondrial chaperone [67] in adult muscle but its role in myoblast differentiation is less well explored, but warrants further investigation based on the current finding that CH60 exhibits similar responses to members of the 70 kDa and 90 kDa heat shock protein families.

The enzyme, 6PGD, was unique in that it was the only protein amongst those investigated that exhibited a significant gain in abundance during the later period of differentiation specifically in aged cells. 6PGD is the rate limiting enzyme in the pentose phosphate pathway [68] and catalyses the reaction between ribulose 5-phosphate and 6-phosphogluconate, which produces NADPH [69] that may be associated with a heightened generation of reactive oxygen species (ROS) [70]. In C_2_C_12_ myoblasts and myotubes, increased ROS via myostatin treatment upregulated the activity of NADPH oxidases via the translocation and activation of NF-Kβ to the nucleus [71]. Induction of ROS activated TNF-α which upregulated NF-Kβ signalling and acted as a feedforward loop to increase secretion of myostatin [71], further accelerated the induction of these signalling pathways. However, based on the current data it is unclear whether the rise in 6PGD abundance (and potentially ROS production) further accelerates damage or, alternatively, represents a counter attempt to recover losses in proteostasis in aged cells. For example, elevated G6PD abundance could promote cell survival by protecting against elevated ROS producton via elevated TNF-α activation of NF-Kβ. While TNF-α activation of NF-Kβ were not investigated in the current work, this response would be expected to lead to lesser expression of MyoD [12]. Moreover, T cells with genetically reduced G6PD content display impaired cell function as a result of reduced cytokine production and glycolysis [72]. In the same study, NF-Kβ inducing kinase (NIK) directly phosphorylated G6PD and overexpression of NIK subsequently elevated G6PD. Therefore, elevated G6PD could act to protect cells from excessive ROS production (potentially activated by elevated myostatin) through elevated NF-Kβ and NIK. Consequently, elevated NF-Kβ which is activated via TNF-α suppresses MyoD and greater TNF-α and decreased MyoD were reported in replicatively aged cells [12].

Consistent with our earlier work [12], the expression of IGF-I was suppressed in replicatively aged myoblasts, which may be expected to have a further negative impact on Akt-mTOR signalling and subsequently differentiation and hypertrophy [24]. In addition, myostatin expression was elevated and may further inhibit IGF-I stimulated growth pathways [73] possibly through secreted myostatin [74]. Indeed, myostatin treatment on human muscle cells reduces Akt–mTOR and inhibits differentiation and myotube size via reduction in MyoD and myogenin expression [26]. Myostatin also decreased the expression of E3 ligases MuRF1 and MAFbx in muscle cells [26]. This study supports our findings that demonstrate elevated myostatin expression and reduced Akt-mTOR signalling at 0 h and 24 h as well as impaired degradation. Moreover, genetically altered myostatin peptides that elevated myostatin expression inhibited C_2_C_12_ myoblasts and myotube mixed protein synthesis although there was no difference in estimated degradation rates [75]. In addition, in Japanese Flounder *Paralichthys olivaceus* muscle cells, overexpression of myostatin attenuated all of the MRFs and Akt and mTOR signalling and induced FOXO, MuRF1 and MAFbx expression [76]. Proteo-ADPT assesses the degradation rate directly on a protein-by-protein basis and future more in-depth studies could offer new insight regarding such E3 ligases target specific proteins for degradation, (e.g. cytoskeletal proteins [56]) and may not equate to the overall degradation rate of all proteins in a particular cell.

Glycolysis is the dominant energy source in proliferating myoblasts and upon differentiation there is a gradual transition to a greater contribution of ATP resynthesis by oxidative phosphorylation [77]. Seven out of eight enzymes of glycolytic metabolism exhibited significant disruption in either abundance or turnover in aged compared to young cells. Proteo-ADPT data for glycolytic enzymes were markedly different to the patterns exhibited by ribosomal and chaperone proteins. In young cells, there was a general decline in the abundance of glycolytic enzymes, which is consistent with our earlier findings and the greater energy requirements associated with higher levels of protein synthesis in myoblasts compared to myotubes [28]. In aged myoblasts, 4 glycolytic proteins exhibiting increased abundance and the turnover of 7 glycolytic enzymes was impaired in aged compared to young myoblasts during early differentiation. Dysfunction of glycolytic energy metabolism is associated with an accelerated onset of brain aging and disorders such as Alzheimer’s [80]. For instance, TPIS, the mid-point catalyst between hexose and triose stages of glycolysis, exhibited impaired turnover and no difference in the rate of abundance change in aged compared to young myoblasts. In support, TPIS protein content was impaired in the aged brain and consequently, there was reduced NADH metabolism and an increase in the bioproduct methylglycoxal which activates advanced glycation end-products [81].

Our findings point to age effect on UPS or other degradation processes, but protein subunits of UPS and autophagy were not specifically identified within the current analyses. Further work using higher resolution MS is warranted in this model of muscle ageing. In *C elegans* turnover and abundance of proteasome subunits is maintained during ageing but muscle proteins are not particularly enriched in whole worms. Ubiquitinated proteins accumulate in liver of aged mice [39] suggesting impaired degradation occurred earlier in life supporting the role of UPS in healthy ageing. Future studies should also include time periods that were absent from the current work. It will be important to investigate proteome dynamics in proliferating myoblasts as the current findings support our earlier report [12] that the transition from proliferation to differentiation in replicatively aged C_2_C_12_ lags behind the normal timeline in young cells and ultimately the processes of fusion and myotube hypertrophy fail in replicatively aged cells. Sharples *et al.* [12] reported aged cells continue to undergo proliferation in the minutes immediately after transfer to differentiation media, with a significantly greater proportion of replicatively aged cells in S/G2 phases of the cell cycle and fewer cells in G1 phase in comparison to controls. Suggesting a continued proliferation and inability of replicatively aged cells to exit the cell cycle in G1 and undergo differentiation. Our current analyses further highlight the mid-differentiation period (24 h - 72 h) as a key time where differences in fusion occur and Proteo-ADPT analysis including this timespan may shed new light on the mechanisms that impair fusion of replicatively aged myoblasts.

In conclusion, replicatively aged myoblasts exhibit significant disruption in the normal proteome dynamics associated with differentiation in young cells (Figure 8). In particular, our data suggest the quality of key chaperone proteins, including HS90A, HS90B and HSP7C were reduced in aged cells and may account for the disruption to cell signalling required for differentiation, fusion and myotube growth. Our findings and the associated literature suggest a failure in protein degradation may cause disruption to ribosomal capacity that subsequently impairs protein synthesis, and in muscle may contribute to the development of sarcopenia.

## Declarations

## Acknowledgements

Authors would like to thank Liverpool John Moores University for providing funding for the project.

## Conflict of interest statement

The authors declare no conflicts of interest.

## Author contributions

A.D.B designed the study, performed experiments and data analyses, wrote and edited the manuscript. C.E.S designed the study and edited the manuscript. J.G.B designed the study, performed data analysis and edited the manuscript.

## Data Availability

Raw MS/MS data sets are provided in the supplementary documents, any other data is available from the corresponding author upon reasonable request.

## Notes

### Competing Interest Statement

The authors have declared no competing interest.

