## Supplementary tables and figures for "Degradation of ribosome and chaperone proteins is attenuated during the Differentiation of Replicatively Aged C2C12 Myoblasts"

**Table S1. Primer Sequences and Product Lengths.**

| **Gene** | **Accession** | **Sequence**  **(Forward/Reverse)** | **Product**  **Length (bp)** |
| --- | --- | --- | --- |
| RP2β  (aka polr2b) | NM_153798.2 | F: GATCCGTATCAGGTTCATGTCTGTA  R: CGGCATTCGCACCAAGGAA | 123 |
| Myogenin | NM_031189 | F: CAACCCAGGAGATCATTTGCT  R: TGACAGACAATCTCAGTTGGG | 145 |
| Myostatin | NM_010834 | F: GACCCGTCAAGACTCCTACA  R: CTTCACATCAATACTCTGCCAAATA | 146 |
| IGF-I | NM_010512 | F: CATCGTCTTTAGGAGTGTTTGTTT  R: TAGAATGTTATTGATAAAGGACCAAC | 98 |
| IGF-II | NM_010514.3 | F: CTCCTGGAGACATACTGTGCC  R: AGGTGTCATATTGGAAGAACTTGC | 121 |

**Table S2. Gene Basal C_T_ Values in Young and Aged**

| **Gene** | **Basal Young**  **C _T_ Values** | **Basal Aged**  **C _T_ Values** | **Significance (basal C _T_ Y vs O)**  **(P < 0.05)** |
| --- | --- | --- | --- |
| RP2β (Polr2b) | 23.56±1.38 | 23.47±1.29 | NS |
| Myogenin | 21.82±0.73 | 32.19±1.43 | P < 0.001 |
| Myostatin | 34.90±0.21 | 29.14±0.41 | P < 0.001 |
| IGF-I | 28.10±0.57 | 31.93±0.82 | P < 0.001 |
| IGF-II | 26.71±1.08 | 32.10±1.54 | P < 0.001 |

**Table S3. Protein Concentrations Quantified by BCA Assay**

|  |  | **Young** | **Aged** |
| --- | --- | --- | --- |
| **Cell** | **Time** | **Average ± SD (μg/well)** | **Average ± SD (μg/well)** |
| Myoblast | 0h | 404.55 ± 17.66 | 334.24 ± 8.14 |
|  | 24h | 459.55 ± 36.36 | 487.77 ± 33.27 |
| Myotube | 72h | 452.38 ± 15.49 | 335.92 ± 24.06 |
|  | 96h | 512.86 ± 34.60 | 324.95 ± 3.45 |

**Table S4. Abundance, synthesis and degradation of proteins associated with glycolysis and pyruvate metabolism in young and replicatively aged myoblasts during early differentiation.**

| **Accession** | **Age** | **ABD 0h (ng/well)** | **ABR (ng/well/h)** | **ASR (ng/well/h)** | **ADR (ng/well/h)** | **ABD 24h (ng/well)** |
| --- | --- | --- | --- | --- | --- | --- |
| **Glycolysis and Pyruvate Metabolism** | | | | | | |
| ALDOA | Y | 799.11 ± 4.56 | 7.47 ± 0.2 | 50.57 ± 0.53 | 43.1 ± 0.44 | 978.36 ± 9.27 |
|  | A | 759.92 ± 3.28 | 15.26 ± 0.3 | 38.22 ± 0.37 | 22.96 ± 0.62 | 1126.2 ± 5.46 |
| ENOA | Y | 1337.9 ± 3.14 | 19.96 ± 0.59 | 80.82 ± 0.6 | 60.85 ± 0.35 | 1817.06 ± 16.95 |
|  | A | 1280.91 ± 4.72 | 31.96 ± 0.27 | 63.32 ± 0.13 | 31.36 ± 0.28 | 2047.97 ± 6.72 |
| G3P | Y | 1773.37 ± 13.66 | 8.84 ± 0.29 | 97.77 ± 1.07 | 88.92 ± 0.99 | 1985.57 ± 16.1 |
|  | A | 1537.14 ± 7.15 | 29.44 ± 0.42 | 82.42 ± 0.23 | 52.98 ± 0.28 | 2243.71 ± 4 |
| KPYM | Y | 1795.54 ± 10.5 | 9.76 ± 0.47 | 96.69 ± 1.19 | 86.93 ± 0.77 | 2029.81 ± 18.01 |
|  | A | 1614.84 ± 6.06 | 21.63 ± 0.66 | 64.96 ± 0.29 | 43.33 ± 0.9 | 2134.04 ± 13.61 |
| LDHA | Y | 579.69 ± 4.55 | 4.12 ± 0.08 | 33.07 ± 0.2 | 28.95 ± 0.24 | 678.48 ± 3.84 |
|  | A | 582.90 ± 5.00 | 7.88 ± 0.38 | 23.77 ± 0.33 | 15.89 ± 0.68 | 771.97 ± 5.32 |
| PGAM1 | Y | 231.97 ± 1.48 | 0.66 ± 0.03 | 10.88 ± 0.28 | 10.22 ± 0.26 | 247.77 ± 1.19 |
|  | A | 236.66 ± 0.91 | 2.99 ± 0.09 | 9.24 ± 0.14 | 6.25 ± 0.11 | 308.42 ± 1.8 |
| PGK1 | Y | 605.94 ± 2.36 | 5.06 ± 0.27 | 42.99 ± 0.39 | 37.93 ± 0.4 | 727.34 ± 6.62 |
|  | A | 578.91 ± 3.49 | 9.15 ± 0.04 | 26.89 ± 0.56 | 17.75 ± 0.56 | 798.43 ± 3.37 |
| TPIS | Y | 242.59 ± 1.48 | 1.54 ± 0.05 | 13.98 ± 0.02 | 12.44 ± 0.04 | 279.54 ± 1.74 |
|  | A | 244.19 ± 1.96 | 4.05 ± 0.14 | 8.32 ± 0.15 | 4.26 ± 0.21 | 341.48 ± 1.74 |

**Table S5. Abundance, synthesis and degradation of proteins associated with translation in young and replicatively aged myoblasts during early differentiation.**

| **Accession** | **Age** | **ABD 0h (ng/well)** | **ABR (ng/well/h)** | **ASR (ng/well/h)** | **ADR (ng/well/h)** | **ABD 24h (ng/well)** |
| --- | --- | --- | --- | --- | --- | --- |
| **Translation Elongation** | | | | | | |
| EF1A1 | Y | 1983.52 ± 13.58 | 6.01 ± 0.3 | 91.73 ± 0.5 | 85.71 ± 0.64 | 2127.86 ± 19.24 |
|  | A | 1458.55 ± 3.39 | 29.53 ± 0.64 | 78.43 ± 0.34 | 48.91 ± 0.73 | 2167.18 ± 17.3 |
| EF2 | Y | 1098.62 ± 3.26 | 4.24 ± 0.15 | 54.09 ± 0.24 | 49.85 ± 0.12 | 1200.44 ± 4.99 |
|  | A | 866.46 ± 4.82 | 12.97 ± 0.13 | 55.03 ± 0.36 | 42.06 ± 0.34 | 1177.77 ± 6.64 |
| IF4A1 | Y | 199.24 ± 0.57 | 0.93 ± 0.05 | 6.08 ± 0.09 | 5.15 ± 0.14 | 221.64 ± 1.79 |
|  | A | 144.08 ± 0.38 | 2.66 ± 0.03 | 4.46 ± 0.03 | 1.8 ± 0.01 | 207.92 ± 0.71 |
| **Large Ribosomal Subunit** | | | | | | |
| RL11 | Y | 97.82 ± 0.61 | -0.04 ± 0.04 | 3.97 ± 0.11 | 4.01 ± 0.13 | 96.94 ± 0.55 |
|  | A | 60.44 ± 0.15 | 0.94 ± 0.01 | 0.82 ± 0.02 | -0.12 ± 0.02 | 82.99 ± 0.43 |
| RL19 | Y | 132.24 ± 1.18 | 0.05 ± 0.06 | 8.16 ± 0.02 | 8.11 ± 0.07 | 133.35 ± 1.11 |
|  | A | 85.54 ± 0.13 | 1.64 ± 0.04 | 5.05 ± 0.13 | 3.41 ± 0.12 | 124.8 ± 1.05 |
| RL22 | Y | 244.26 ± 1.24 | 1.06 ± 0.05 | 19.41 ± 0.09 | 18.35 ± 0.14 | 269.68 ± 1.75 |
|  | A | 187.29 ± 1.31 | 2.96 ± 0.03 | 11.74 ± 0.18 | 8.78 ± 0.18 | 258.34 ± 1.82 |
| RL27A | Y | 80.81 ± 0.16 | 0 ± 0.04 | 2.77 ± 0.02 | 2.77 ± 0.05 | 80.92 ± 1.03 |
|  | A | 50.28 ± 0.57 | 0.76 ± 0.01 | 1.62 ± 0.1 | 0.87 ± 0.11 | 68.42 ± 0.32 |
| RL30 | Y | 77.64 ± 0.36 | -0.36 ± 0.02 | 1.37 ± 0.06 | 1.73 ± 0.07 | 69.04 ± 0.64 |
|  | A | 52.6 ± 0.31 | 0.62 ± 0.01 | 1.6 ± 0.02 | 0.98 ± 0.03 | 67.45 ± 0.61 |
| RL35 | Y | 219.85 ± 1.25 | 0.26 ± 0.07 | 2.29 ± 0.17 | 2.04 ± 0.16 | 226.04 ± 2.3 |
|  | A | 173.23 ± 1.33 | 2.65 ± 0.05 | 5.03 ± 0.44 | 2.38 ± 0.47 | 236.92 ± 2.07 |
| RL36 | Y | 81.89 ± 0.11 | 0.6 ± 0.03 | 5.42 ± 0.03 | 4.82 ± 0.06 | 96.25 ± 0.87 |
|  | A | 57.37 ± 0.39 | 1.31 ± 0.02 | 7.11 ± 0.1 | 5.8 ± 0.11 | 88.73 ± 0.35 |
| RL7 | Y | 187.82 ± 0.24 | 0.33 ± 0.08 | 8.18 ± 0.11 | 7.85 ± 0.05 | 195.82 ± 1.84 |
|  | A | 121.63 ± 0.98 | 1.82 ± 0.03 | 8.87 ± 0.09 | 7.05 ± 0.06 | 165.31 ± 0.81 |
| RLA2 | Y | 87.67 ± 0.19 | 0.14 ± 0 | 1.51 ± 0.02 | 1.37 ± 0.02 | 91.07 ± 0.31 |
|  | A | 55.54 ± 0.34 | 0.86 ± 0.02 | 2.03 ± 0.02 | 1.17 ± 0.01 | 76.15 ± 0.74 |
| **Small Ribosomal Subunit** | | | | | | |
| RS14 | Y | 136.63 ± 0.98 | 0.43 ± 0.05 | 4.56 ± 0.15 | 4.13 ± 0.19 | 147.03 ± 1.37 |
|  | A | 92.49 ± 0.16 | 2.27 ± 0.03 | 4.05 ± 0.11 | 1.78 ± 0.09 | 146.98 ± 0.94 |
| RS18 | Y | 85.05 ± 0.26 | 0.34 ± 0.04 | 2.15 ± 0.06 | 1.81 ± 0.07 | 93.21 ± 0.67 |
|  | A | 57.16 ± 0.1 | 1.23 ± 0.04 | 1.48 ± 0.09 | 0.24 ± 0.12 | 86.79 ± 0.78 |
| RS25 | Y | 75.23 ± 0.08 | 0.24 ± 0.02 | 2.65 ± 0.03 | 2.41 ± 0.01 | 80.92 ± 0.51 |
|  | A | 51.67 ± 0.35 | 0.95 ± 0.02 | 1.69 ± 0.04 | 0.74 ± 0.06 | 74.41 ± 0.34 |
| RS3 | Y | 111.61 ± 0.38 | 0.27 ± 0.04 | 6.35 ± 0.09 | 6.08 ± 0.11 | 118.14 ± 0.84 |
|  | A | 86.67 ± 0.11 | 1.18 ± 0.03 | 10.11 ± 0.42 | 8.93 ± 0.41 | 115.07 ± 0.61 |
| RS3A | Y | 116.65 ± 0.43 | 0.46 ± 0.03 | 6.7 ± 0.32 | 6.24 ± 0.31 | 127.68 ± 1.12 |
|  | A | 76.34 ± 0.13 | 1.52 ± 0.03 | 14.58 ± 0.17 | 13.06 ± 0.15 | 112.87 ± 0.68 |
| RS6 | Y | 164.95 ± 0.39 | -0.52 ± 0.08 | 6.43 ± 0.03 | 6.95 ± 0.06 | 152.47 ± 1.7 |
|  | A | 114.9 ± 0.6 | 1.18 ± 0.01 | 6.6 ± 0.05 | 5.41 ± 0.04 | 143.3 ± 0.75 |
| RS8 | Y | 148.29 ± 0.45 | 0.18 ± 0.07 | 4.11 ± 0.08 | 3.94 ± 0.02 | 152.49 ± 1.66 |
|  | A | 103 ± 0.18 | 1.6 ± 0.03 | 4.7 ± 0.03 | 3.1 ± 0 | 141.37 ± 0.78 |

**Supplementary Figures**

**A**

100 µm

Scale: 0 µm

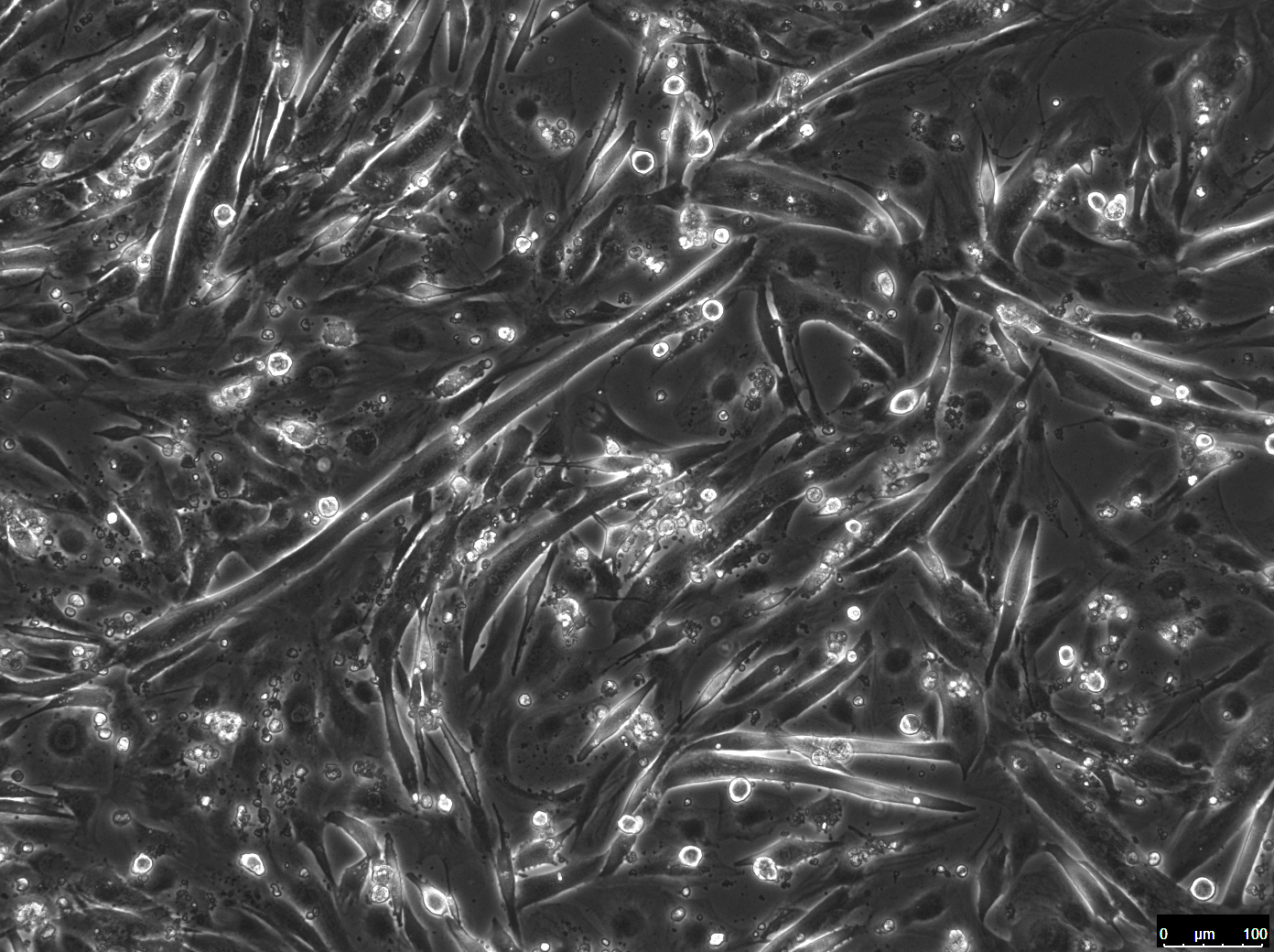

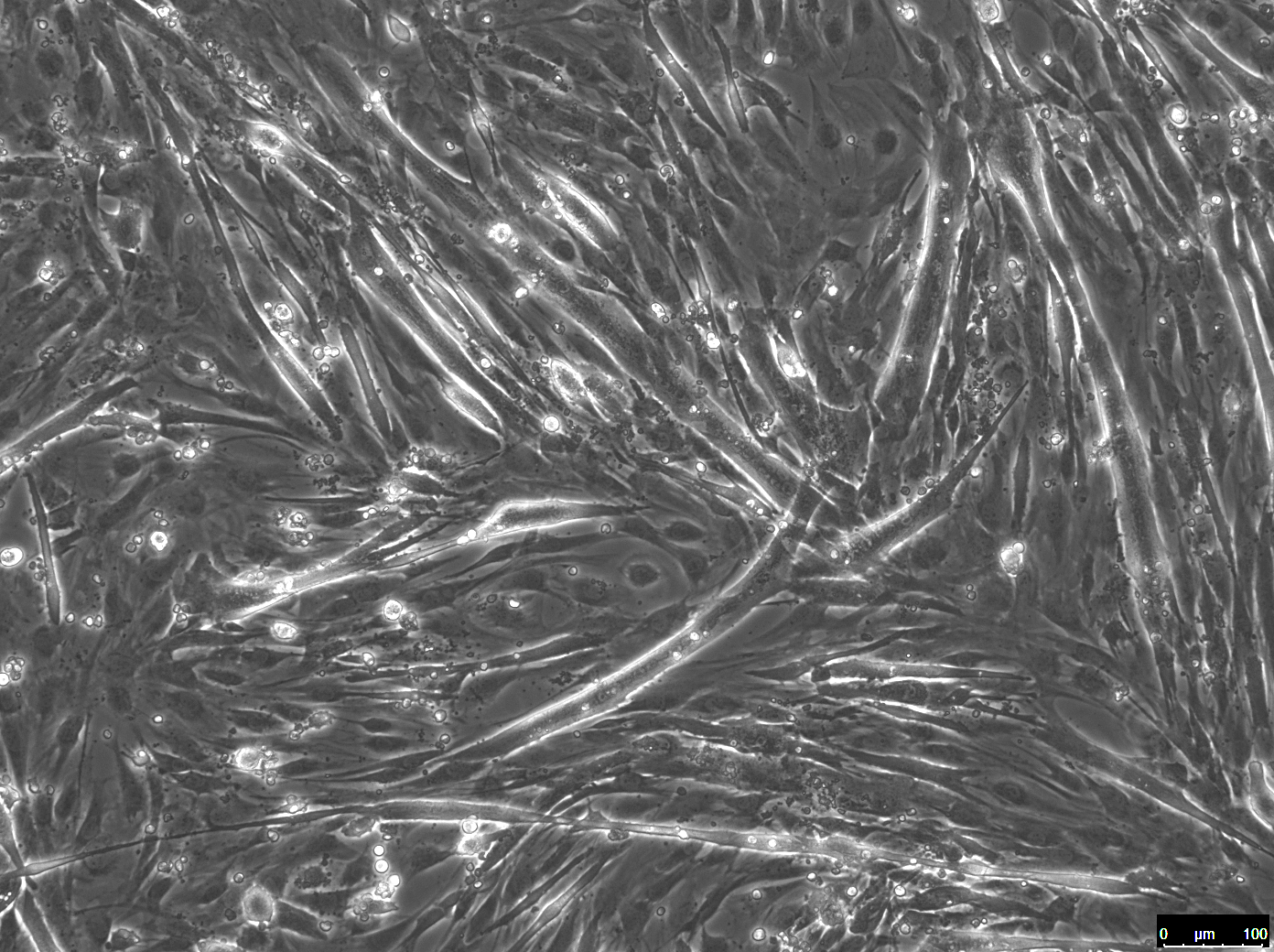

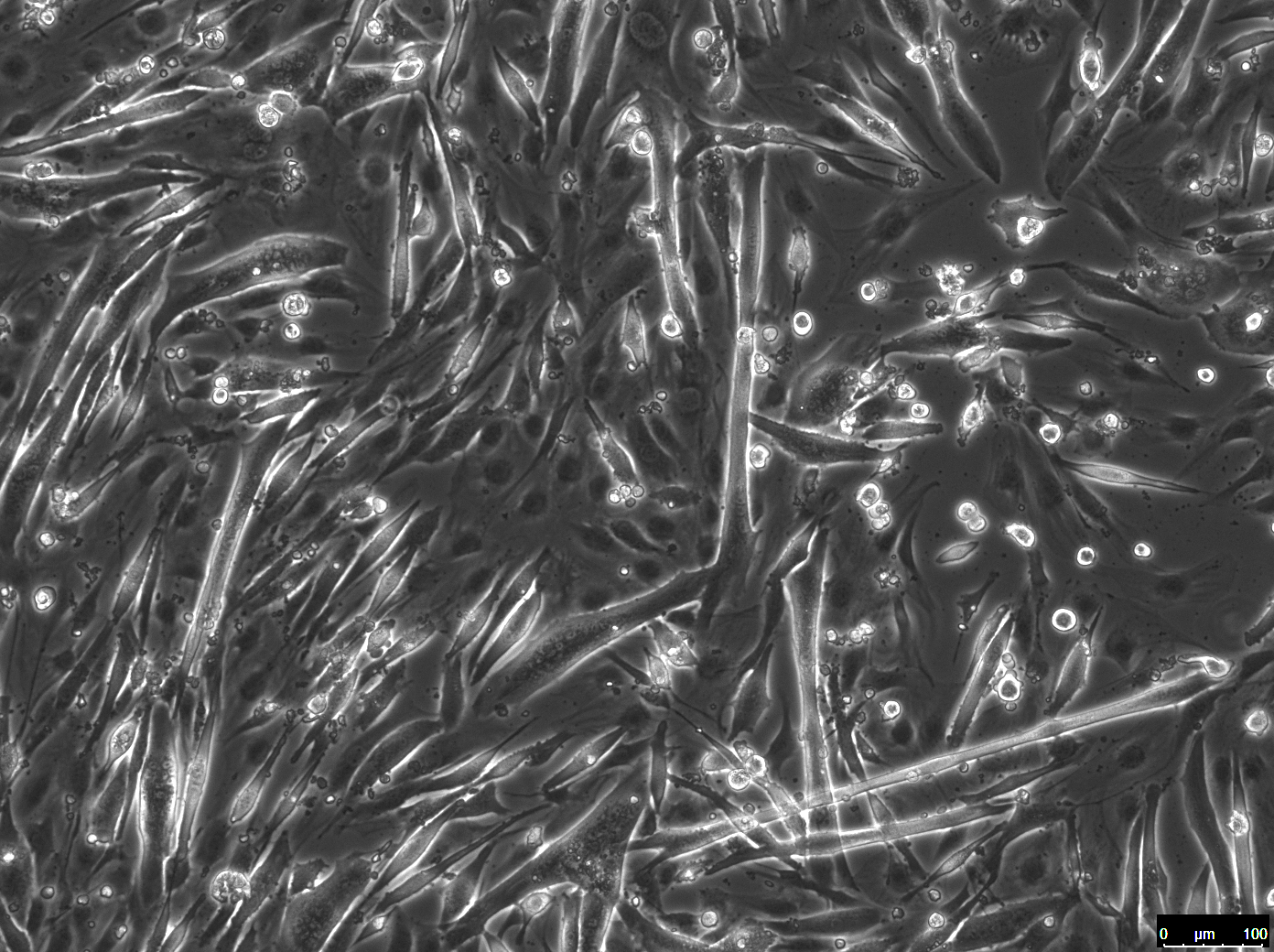

**72h Young DM**

**96h Young DM**

**96h Young DM + 4% D_2_O**

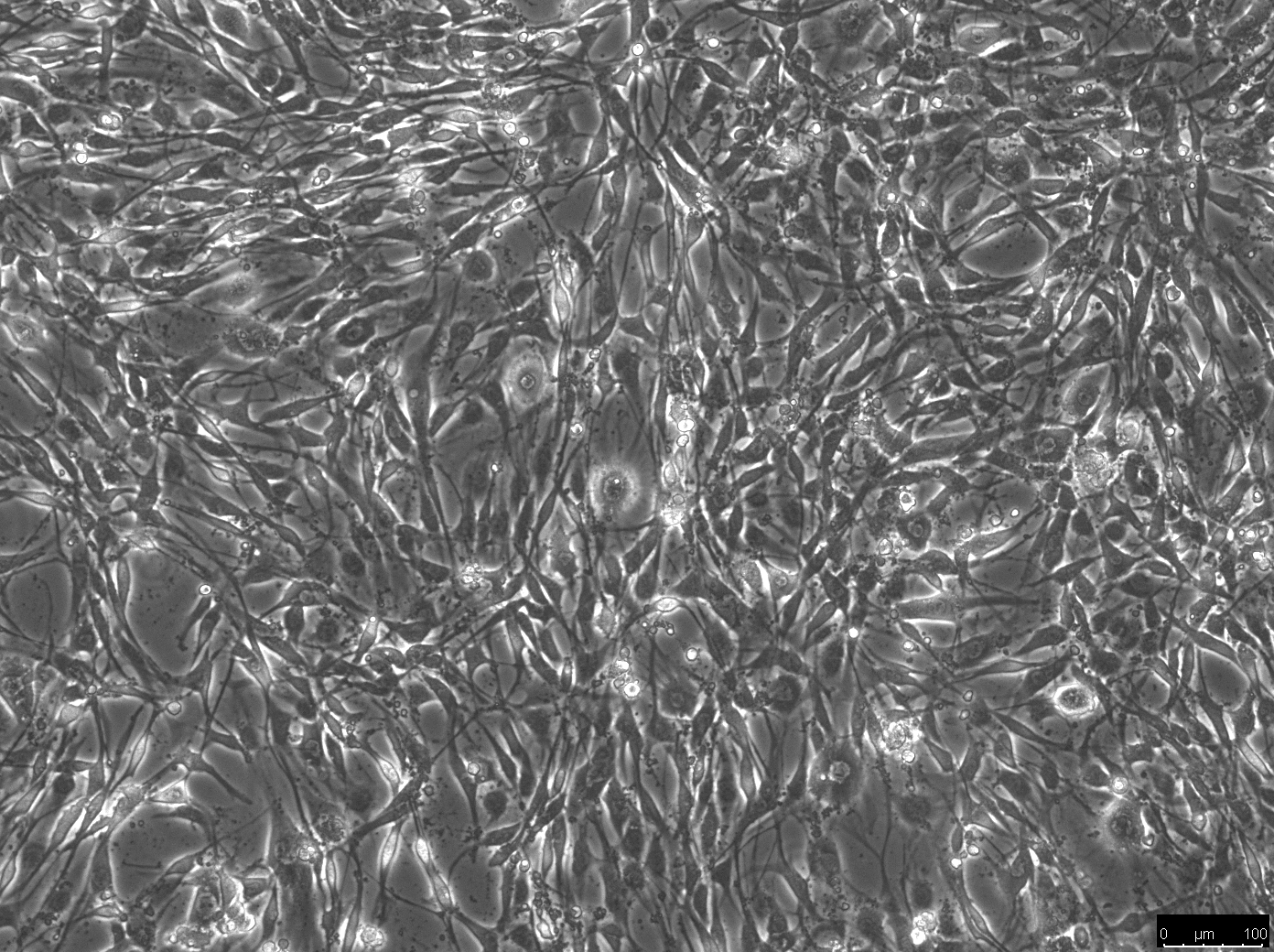

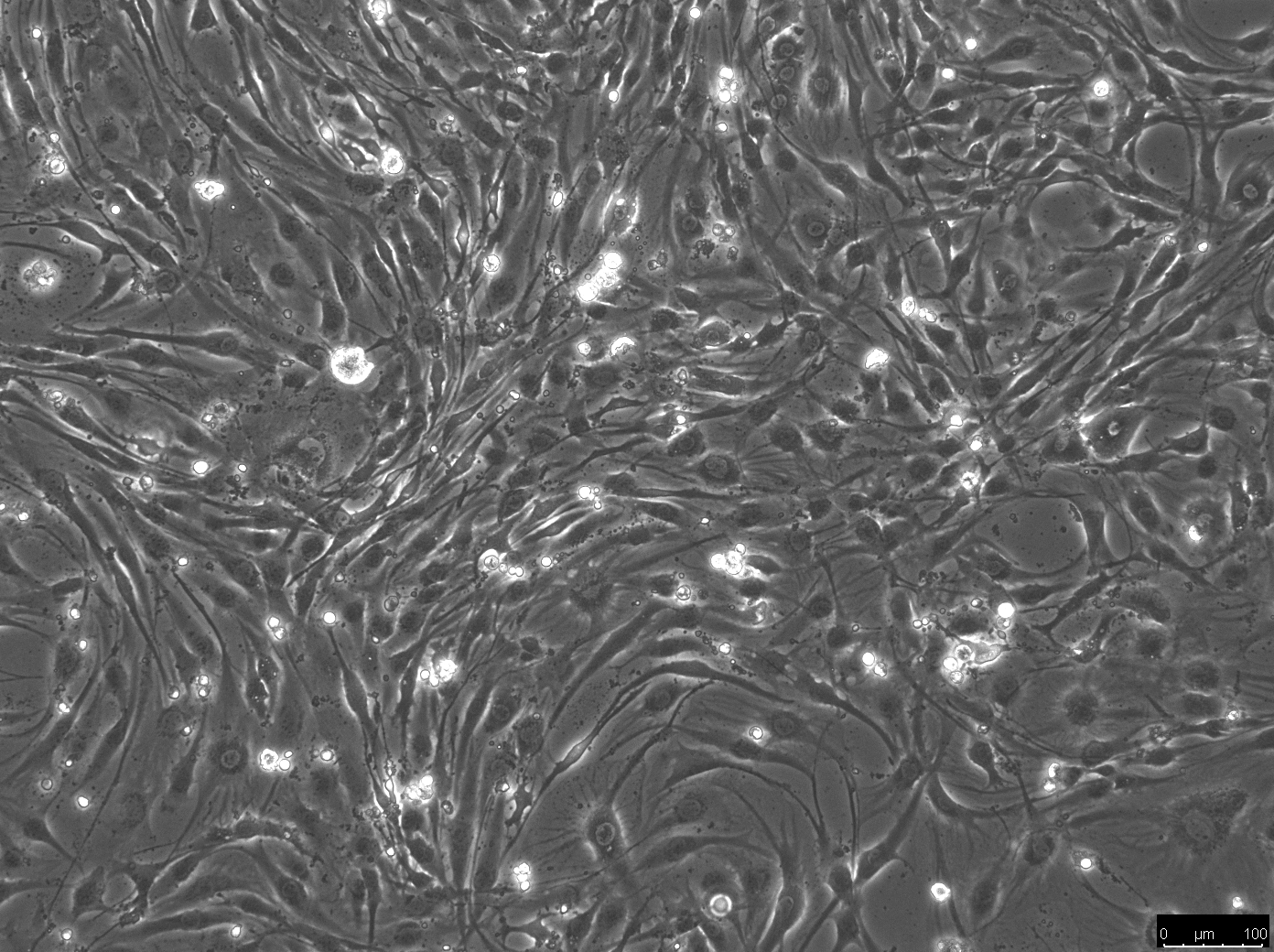

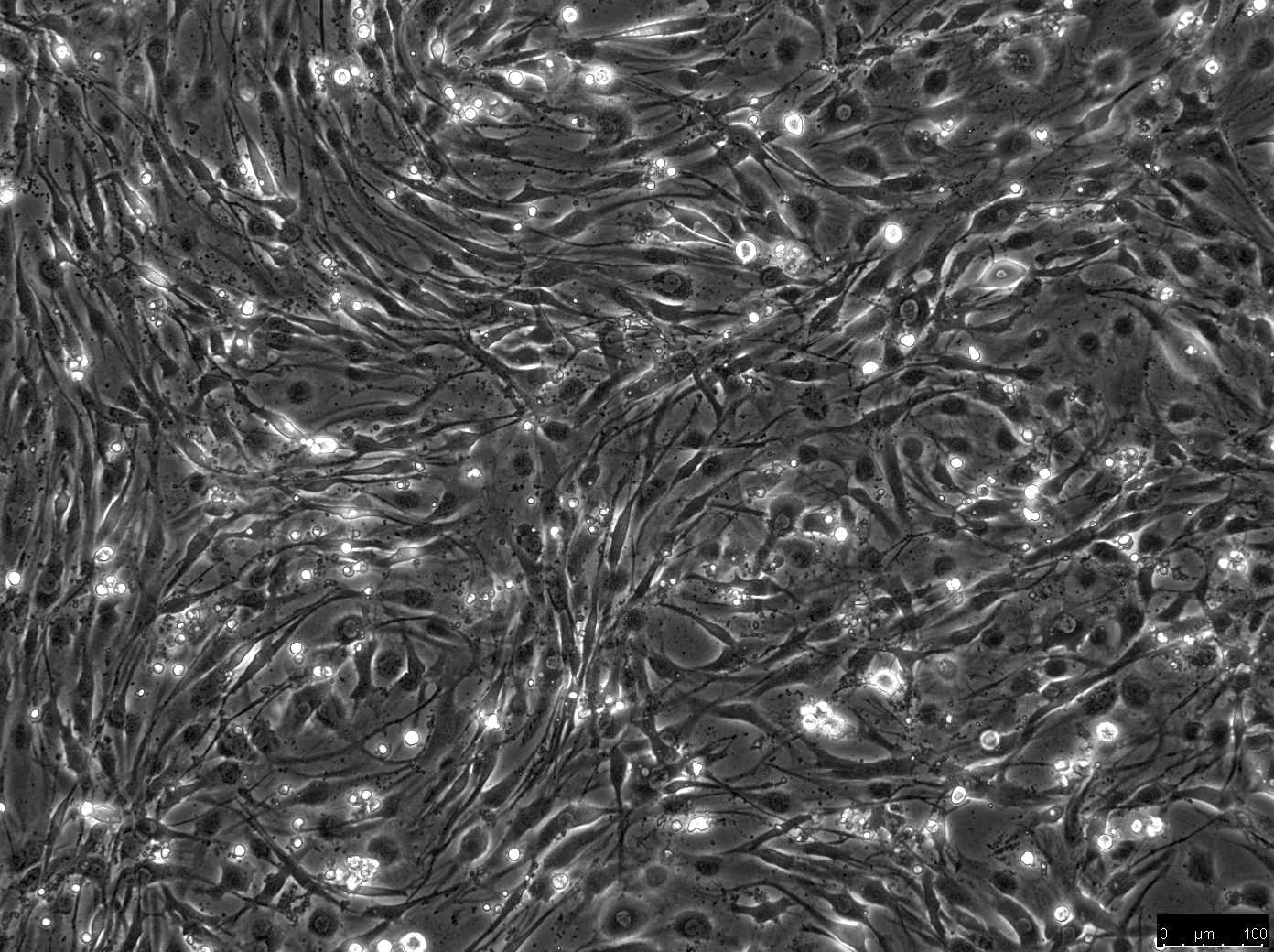

**96h Aged DM**

**72h Aged DM**

**96h Aged DM + 4% D_2_O**

**0h Aged DM**

**24h Aged DM + 4% D_2_O**

**24h Aged DM**

100 µm

Scale: 0 µm

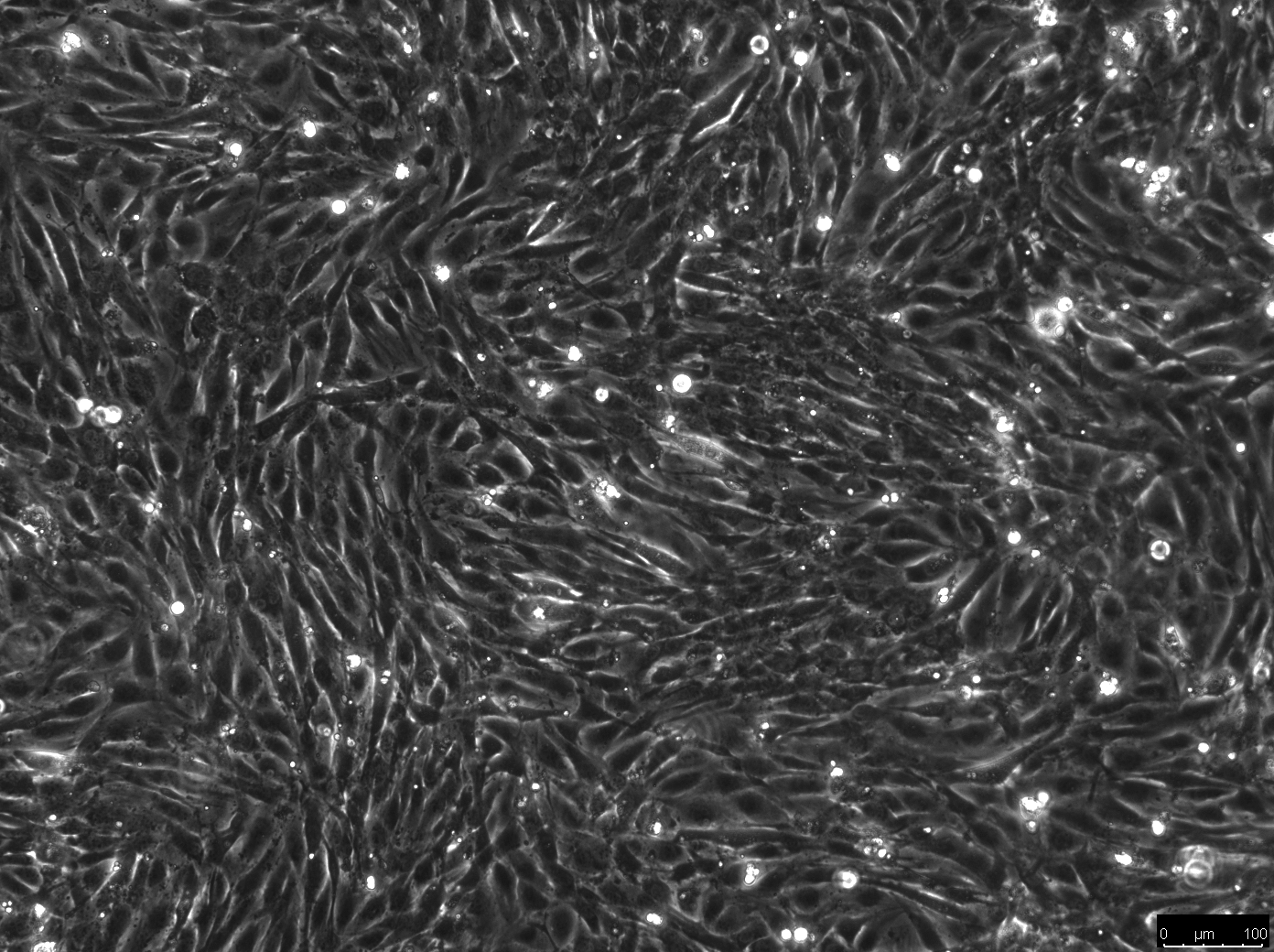

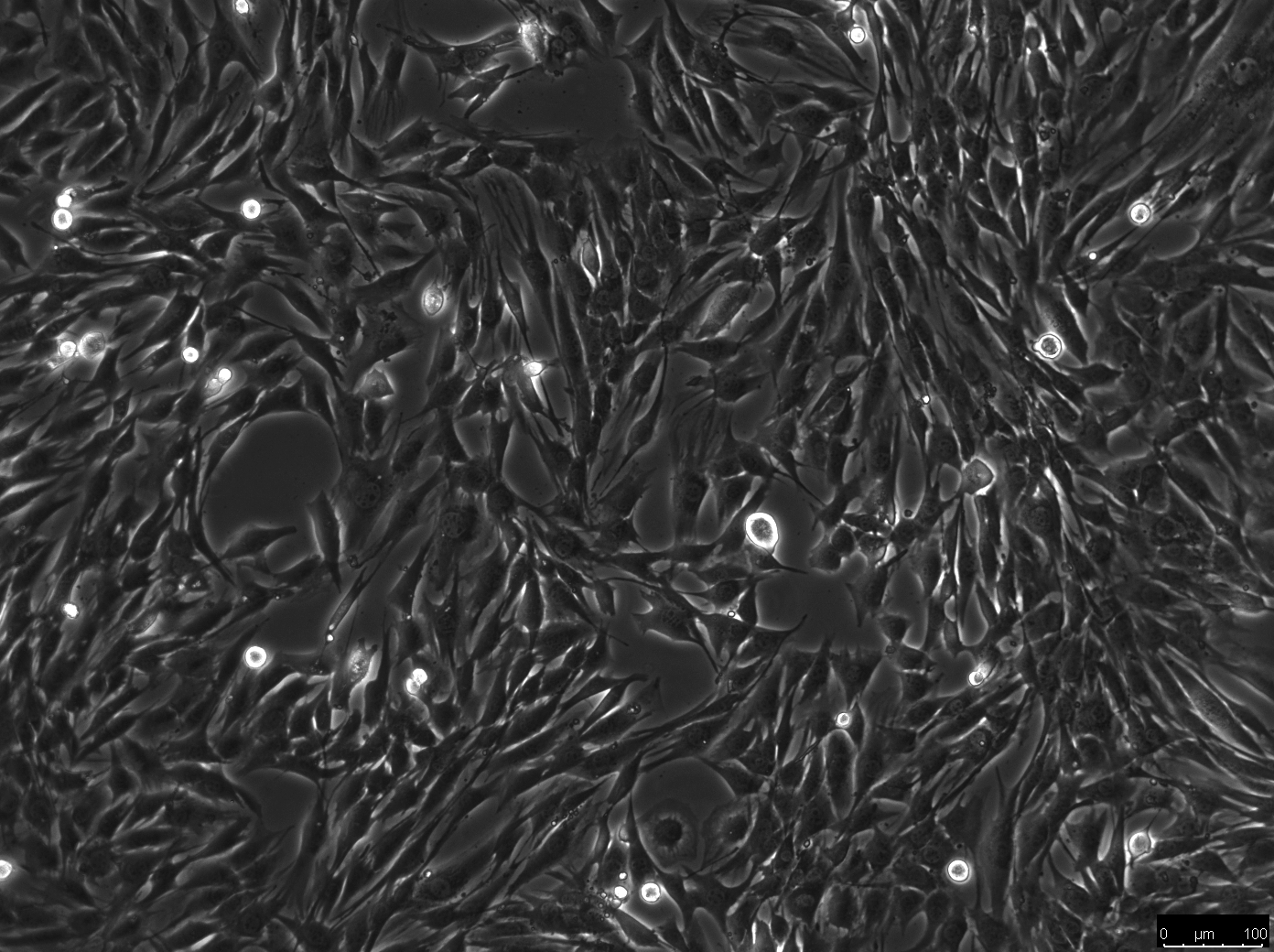

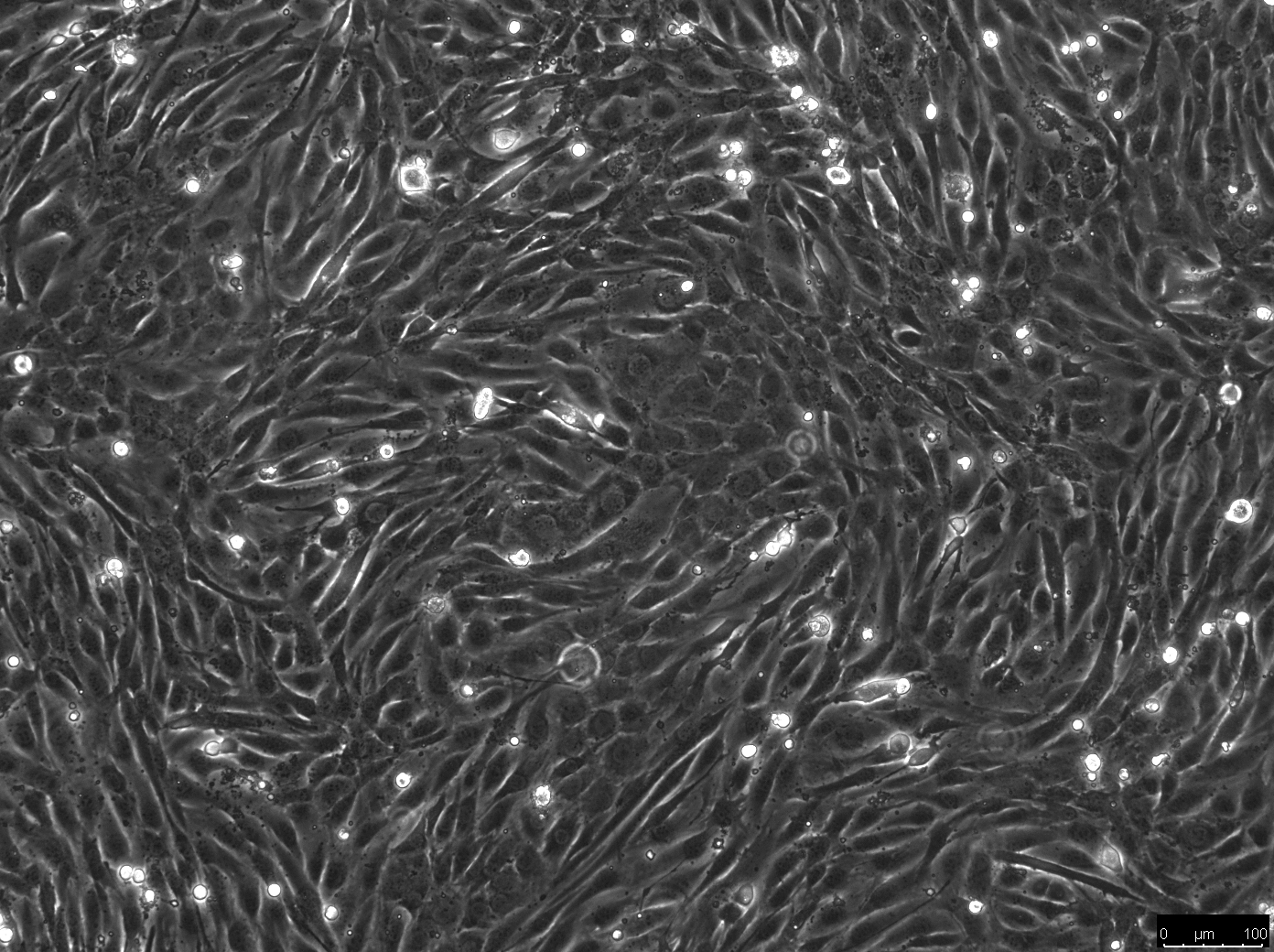

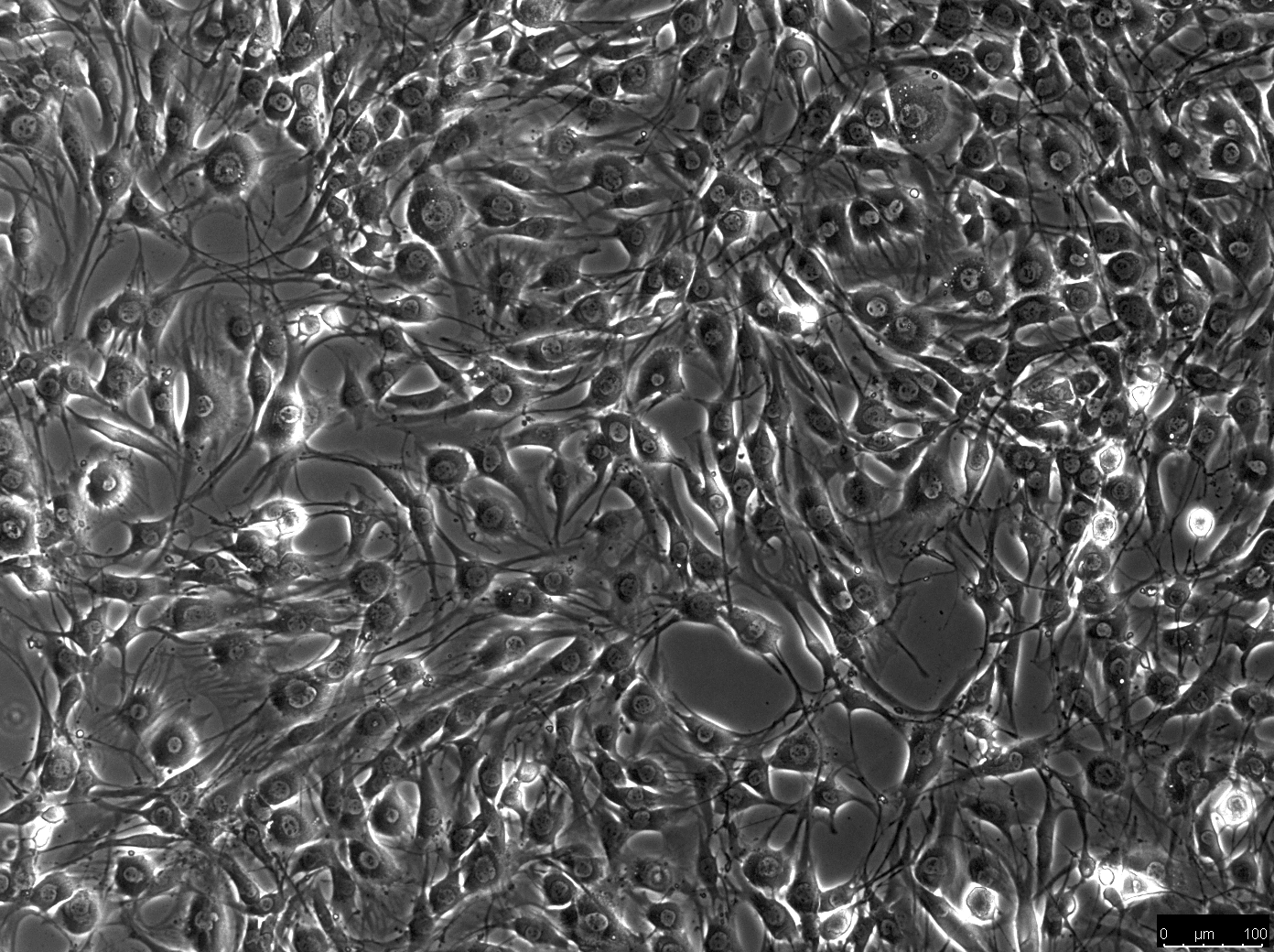

**0h Young DM**

**24h Young DM**

**24h Young DM + 4% D_2_O**

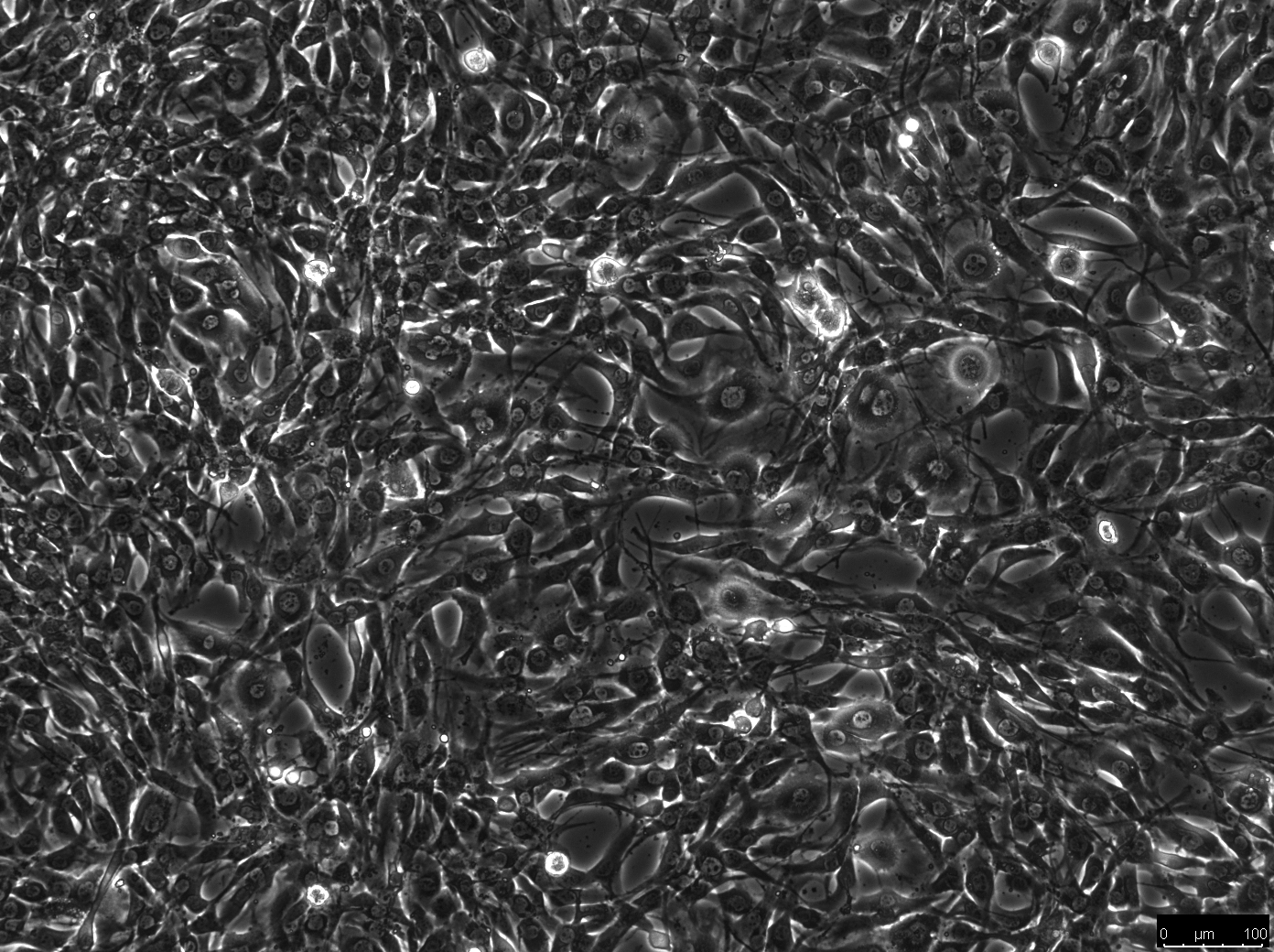

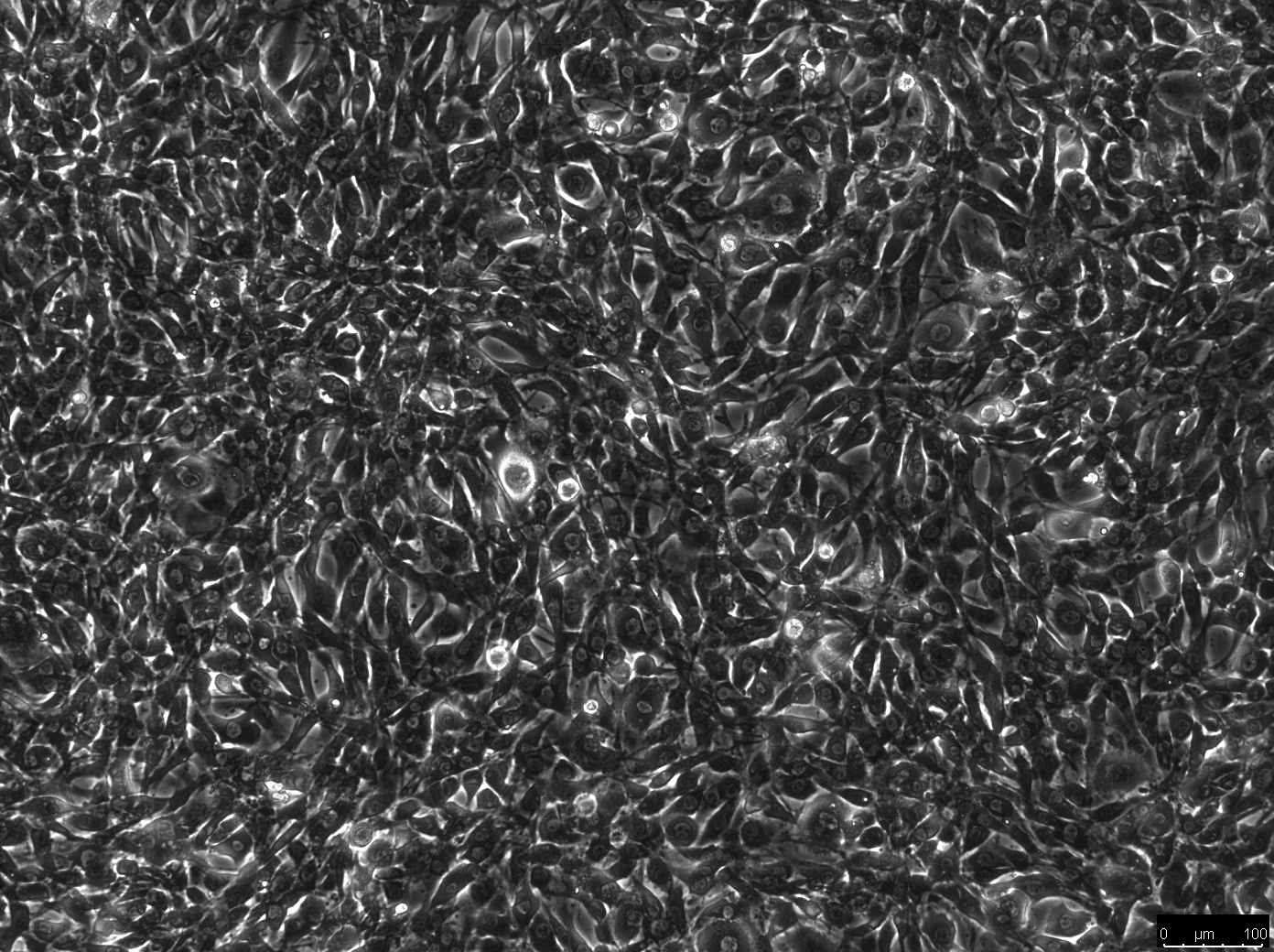

**B**

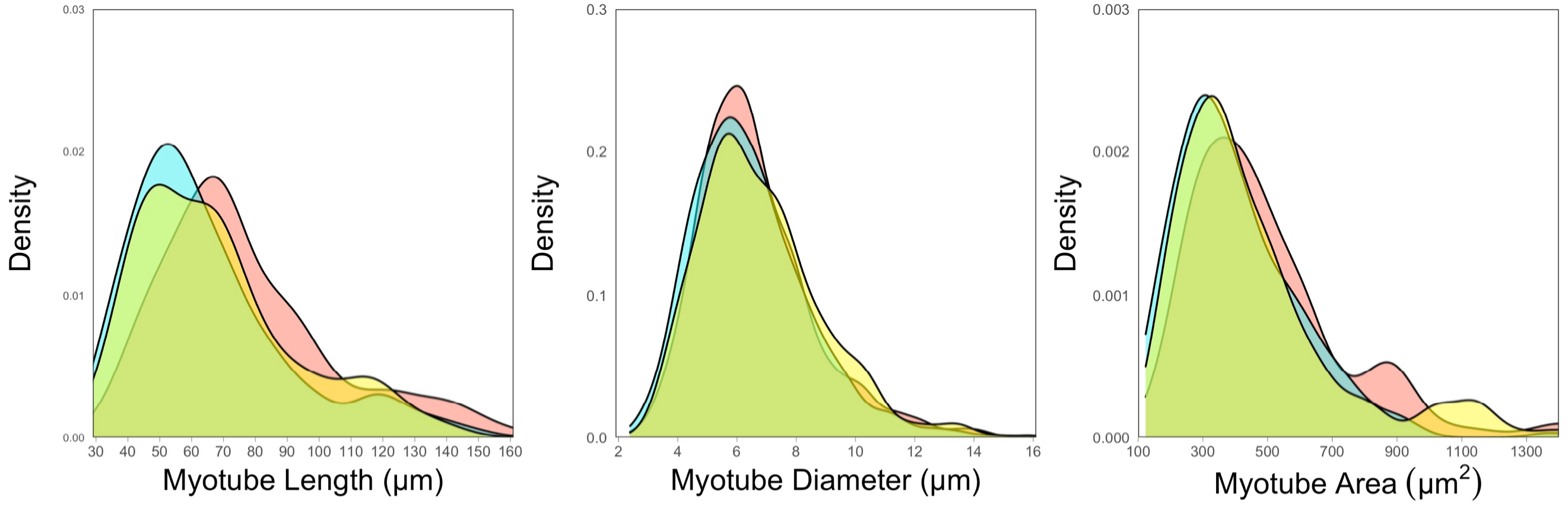

**C**

**Figure S1. Myotube formation was impaired in replicatively aged myoblasts.**

**A/B:** Micrographs in young myoblasts during early (**A**) and later (**B**) differentiation, Scale: 0-100 µm. **C:** Density plots of myotube length (µm), diameter (µm) and area (µm^2^) in young myotubes at 72 h in DM (red) and 96 h in DM (blue) and 4 % D_2_O (yellow). All experiments N = 3 in duplicate.
